# Single-cell informed metabolic modeling reveals organ-specific metabolic adaptations in breast cancer organotropism

**DOI:** 10.64898/2026.09.01.748342

**Authors:** Garhima Arora, Samrat Chatterjee

## Abstract

Breast cancer organotropism is driven by interactions between tumor cells and organ-specific microenvironments that support metastatic growth. To better understand the metabolic basis of organ-specific metastasis, we integrated single-cell transcriptomics with constraint-based systems biology to generate context-specific metabolic models of breast cancer metastasis to the liver, bone, and brain. Our analysis identified both common and organ-specific metabolic changes, suggesting that metastatic cells share a core metabolic program while also adapting to the metabolic environment of each target organ. Primary tumors with metastatic potential showed early alterations in nucleotide metabolism, transport reactions, and energy-related pathways, indicating metabolic changes before metastatic spread. Metabolic transformation analysis identified key metabolic regulators involved in the tricarboxylic acid (TCA) cycle, oxidative phosphorylation, redox balance, and metabolite transport. Integration with CRISPR gene essentiality data further highlighted metabolically important genes as potential therapeutic targets. In addition, analysis of organ-specific secreted metabolites revealed distinct metabolic signatures associated with metastatic colonization of the liver, bone, and brain. Overall, our single-cell-informed metabolic modeling approach shows that breast cancer organotropism is associated with both shared and organ-specific metabolic adaptations. The study provides a framework for identifying potential metabolic vulnerabilities that could be targeted to treat metastatic breast cancer.

## 1 Introduction

Breast cancer is one of the most prevalent cancers and ranks as the second leading cause of cancer-related deaths in women, with an estimated 2.3 million new cancer cases (1 in 4 new cancer cases) and 685,000 cancer deaths (1 in 6 deaths) in 2020 [1–3]. The global landscape projects a substantial increase, with an estimated 2,964,197 new female breast cancer cases anticipated in 2040, signifying a 31% rise from the corresponding figures in 2020 [4]. Research findings suggest that the estimated national costs for medical services and oral prescription drugs related to female breast cancer are expected to rise by 33% in the United States by 2030 [5]. The multifaceted challenge of breast cancer lies in its heterogeneity and propensity for metastasis, a crucial aspect of disease progression. Notably, 20% to 30% of patients diagnosed with early-stage breast cancer will encounter distant metastases, and a staggering 90% of patient fatalities result from complications associated with recurrent or metastatic diseases [6]. The intricacies of distant metastasis unfold through a dynamic and complex multistep process known as the ‘metastatic cascade’. This involves the detachment of tumor cells from the primary tumor, their entry into the systemic circulation (intravasation), survival in the circulation, evasion of immune attacks, adhesion to capillaries, and extravasation before colonizing distant organs [7, 8].

In 1889, Stephen Paget discovered a distinctive pattern in the distribution of organs affected by cancer metastasis, proposing the ‘seed and soil’ hypothesis [9]. According to this hypothesis, cancer cells are referred to as ‘seeds’, and the metastatic destination as the corresponding ‘soil’. The phenomenon, characterized by the nonrandom distribution of metastases, is known as ‘metastatic organotropism’ or ‘organ-specific metastasis’ [10]. Organotropism is governed by a range of factors, including cancer subtypes, molecular attributes of cancer cells, the host’s immune microenvironment, and the interplay with local cells. For example, breast cancer can metastasize to different sites, including bone, lung, liver, and brain; however, different subtypes of breast cancer have different propensities for metastatic sites [11–13]. The spread of cancer cells from the primary tumor to distant sites remains a critical determinant of breast cancer outcomes. Organotropism adds an additional layer of complexity, as metastatic cells exhibit distinct metabolic adaptations that influence their ability to thrive in specific organ microenvironments.

Studies have highlighted the significance of metabolic reprogramming in breast cancer metastasis, emphasizing the role of key metabolic pathways such as glycolysis, oxidative phosphorylation, and lipid metabolism [14–16]. These metabolic alterations are influenced by the molecular subtypes of breast cancer, further shaping the organotropism observed in metastatic spread [17–19]. Moreover, recent evidence suggests that distinct organ microenvironments impose unique metabolic demands on metastatic breast cancer cells, shaping their survival and growth patterns [20–24]. Therefore, it is crucial to understand the intricate connections between metabolism and organotropism in breast cancer for the development of therapeutic strategies. Targeting specific metabolic pathways associated with organ-specific metastasis may offer novel approaches to intervene in the progression of breast cancer.

Recently, a metabolic adaptation mechanism of cancer metastasis has been proposed as an emerging model of the interaction between cancer cells and the host microenvironment, revealing a deep and extensive relationship between cancer metabolism and cancer metastasis. However, research on how the host microenvironment affects cancer metabolism is mostly limited to the local tumour microenvironment at the primary site. There are few studies on how differences between the primary and secondary microenvironments promote metabolic changes during cancer progression or how secondary microenvironments affect cancer cell metastasis preference. Hence, we discuss how cancer cells adapt to and colonize in the metabolic microenvironments of different metastatic sites to establish a metastatic organotropism phenotype. In this study, we aimed to investigate about the tissue-specific metabolic rewiring of metastatic breast cancer cells to resemble the metabolic phenotype of the relevant distal organ during colonization. Genome-scale metabolic models (GSMMs) are computational frameworks that integrate diverse biological data to simulate cellular metabolism at a systems level and have emerged as powerful tools for deciphering metabolic intricacies. Therefore, we employed GSMMs to explore the interplay between cancer cell metabolism and the metabolic niches of distant organs, thereby understanding the molecular determinants governing organ-specific metastasis.

The aim of this study is to investigate the metabolic determinants underlying breast cancer organotropism and to identify the metabolic adaptations acquired by cancer cells during metastatic progression and colonization of secondary organs. By integrating single-cell transcriptomics with genome-scale metabolic modeling, this study seeks to characterize both shared and organ-specific metabolic programs associated with breast metastasis to the liver, bone, and brain. In particular, the study aims to understand how metabolic rewiring in primary tumors contributes to metastatic potential, organ-specific dissemination, and pre-metastatic niche formation, as well as how metastatic cells adapt to distinct microenvironmental constraints at secondary sites.

Understanding both primary and secondary tumors is essential for elucidating the metastatic cascade in breast cancer. Primary tumors harbor early metastasis-enabling traits, including epithelial-to-mesenchymal transition (EMT), immune evasion, dormancy, and metabolic reprogramming, which may influence metastatic fate and organ preference. In contrast, secondary tumors provide insight into the mechanisms by which disseminated tumor cells survive, evade immune surveillance, and adapt to organ-specific conditions such as hypoxia, altered nutrient availability, and oxidative stress. By studying these processes, we would identify key metabolic pathways, genes, and secretory metabolites involved in organ-specific metastasis, thereby providing a foundation for the development of targeted and personalized therapeutic strategies against metastatic breast cancer.

## 2 Results

### 2.1 Single-cell RNA sequence data integration and clustering

For subsequent analysis, scRNA-seq data from 3 patients (see Table 1) including 2 liver metastatic tissues, 1 bone and brain metastatic tissue, and the corresponding 3 primary breast cancer tissues were integrated using canonical correlation analysis (CCA). The process of breast organotropism to different secondary organs is depicted in the Figure 1A. A total of 9151 cells were acquired after quality control and filtering. Using uniform manifold approximation and projection (UMAP), the cluster patterns in the final merged metadata, based on patient and tissue types, were obtained and visualised (see Figures 1B-D). Based on gene expression counts and canonical epithelial cell markers, we observed that almost all markers were expressed across all epithelial cells in each sample, with high percentage expression of genes such as EPCAM, KRT8, and KRT18 (Figure 1E). The distribution of cell counts in primary and metastatic tissues for each patient is shown in Figure 1F. As the disease progressed, the number of epithelial cells decreased. It was in line with our conventional understanding that epithelial-mesenchymal transition (EMT) is a crucial process in tumor progression and metastasis. During EMT, epithelial cells lose their epithelial characteristics and acquire mesenchymal properties, enabling them to become more motile and invasive [25]. The cells undergoing EMT may no longer exhibit typical epithelial markers, contributing to a decrease in the number of epithelial cells in the secondary/metastatic tumors.

**Figure 1:**
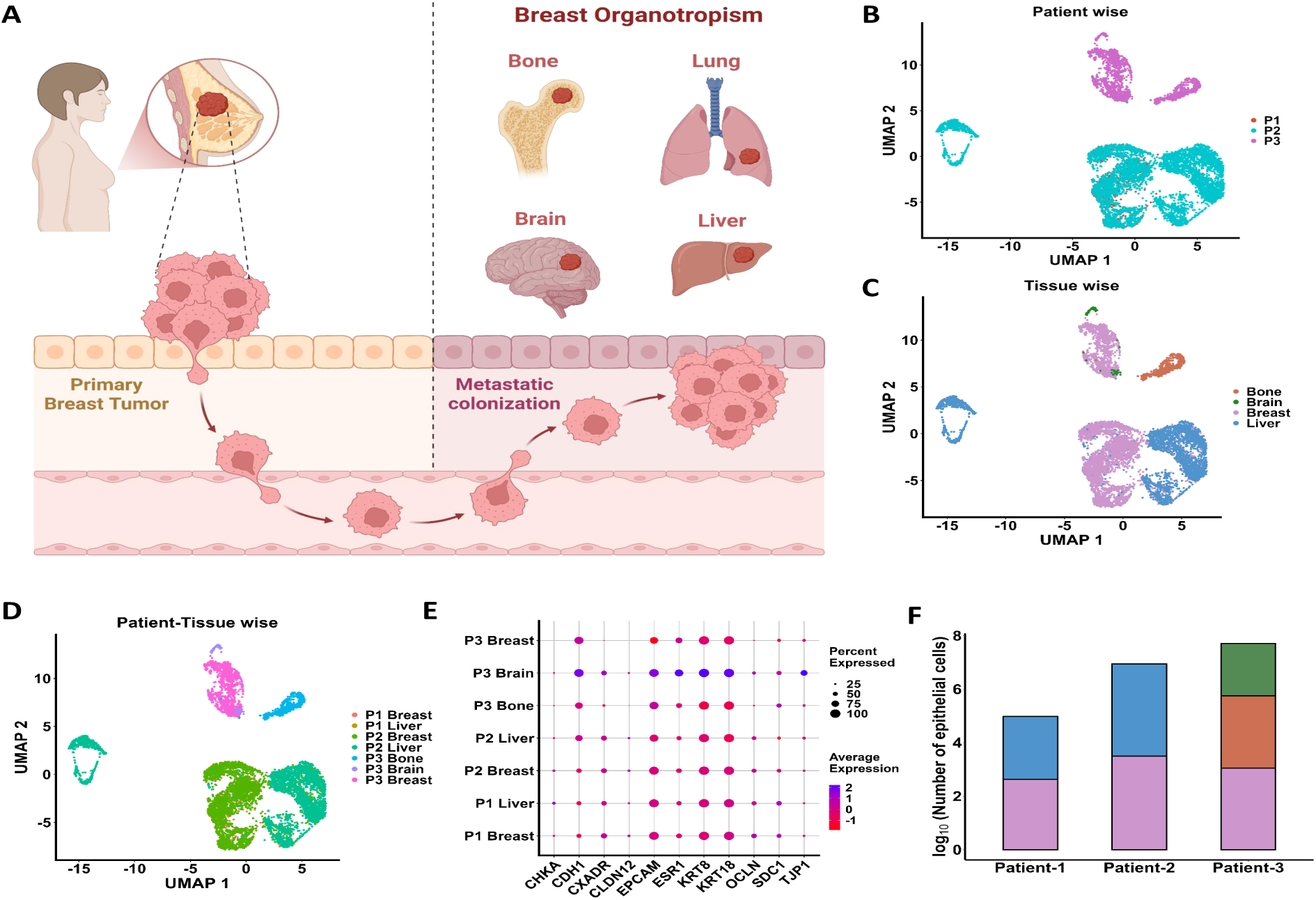
Single-cell atlas of the patients: (A). Breast metastasis to different secondary organs. Uniform manifold approximation and projection (UMAP) of all 9307 cells. Colored by (B). patient type, (C). tissue type, (D). patient-tissue type. P1: patient 1 with liver organotropism; P2: patient 2 with liver organotropism; P3: patient 3 with bone and brain organotropism. (E). The bubble plot shows the expression of epithelial cell markers in each sample. The size of each bubble indicates the percentage of a marker expressed across the epithelial cells of a respective sample, while the colour bar represents their average expression. (F). A stacked bar plot displays the proportion of epithelial cells in primary and metastatic tumor across patients. The distribution of log_10_ of the number of epithelial cells in primary breast tissue is represented by pink color, liver metastatic tissue is represented by blue color, whereas bone and brain metastases are depicted by orange and green color respectively.

**Table 1:** Data description: Number of cells and genes in the data GSE131007.

| Sample name | UCD46 | UCD4 | UCD65 | Combined | Filtered |
| --- | --- | --- | --- | --- | --- |
| **Total cells** | 746 | 6694 | 1867 | 9307 | 9151 |
| Primary tumor | (Breast- 470) | (Breast- 3656) | (Breast- 1159) |  |  |
| Metastatic tumor | (Liver- 276) | (Liver- 3038) | (Bone- 598, Brain- 110) |  |  |
| **Total genes** | 16767 | 19086 | 16816 | 20627 | 20627 |

### 2.2 Generation of organ-specific models

Metabolism is crucial in the multistep complex cascade of tumor dissemination and invasion. Progression as well as metastasis of tumors are accompanied by a significant number of metabolic alterations [26], which present an opportunity for therapeutic targeting of the emerging metabolic liabilities. In this section, we will primarily focus on the organ-specific metabolic adaptations of metastasis. To understand the metabolic differences between primary as well as metastatic tumor, we took single cell RNA sequencing data of three primary breast tumors as well as their corresponding metastatic sites namely, liver, bone and brain. We reconstructed 14 functional metabolic models (see Figure 2) for 7 tissues (one for each cluster) based on the model ‘Human1’ by integrating the expression data, which was carried out by a Task-driven Integrative Network Inference for Tissues (tINIT) algorithm [27–29]. This algorithm ensures the consistency and functionality of the obtained metabolic networks by integrating evidence-based metabolic functions. The cells in each sample were clustered using k-means clustering, and we observed the optimal number of clusters was found to be two in each case (see Figures 3A-G). The data depicting the centroid of each cluster were further used to construct cluster-and tissue-specific models. We observed that all the built models (for model parameters, see Figure 3H) were able to perform all 57 metabolic tasks supplied during model building, ensuring their metabolic functionalities (see Table S1).

**Figure 2:**
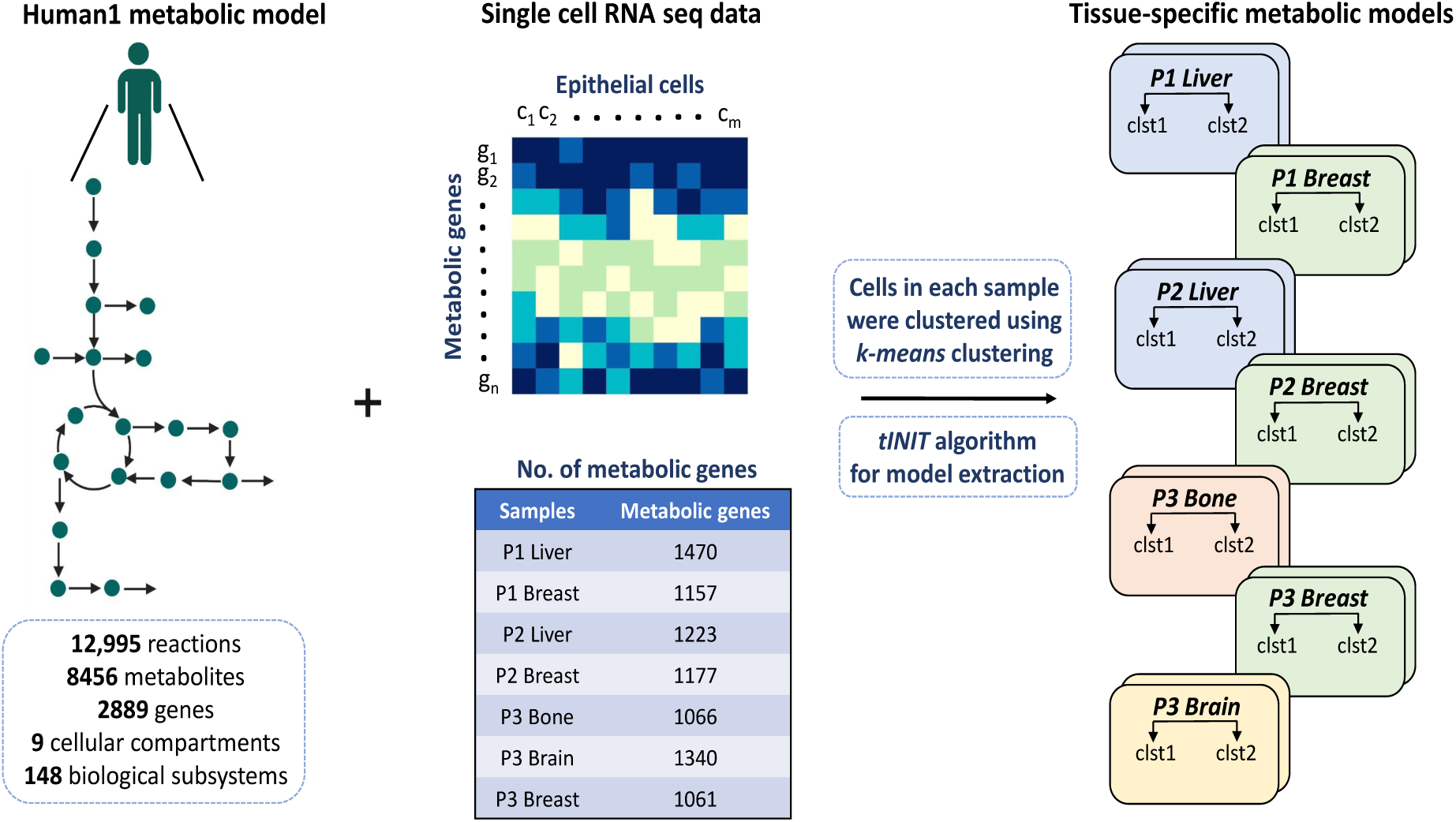
Context-specific metabolic model construction: The figure shows the workflow for constructing tissue-specific metabolic models. The k-means clustering method was used to cluster cells in each sample, and then the single RNA seq data of epithelial cells was integrated into the generic human metabolic model using the tINIT algorithm. Here, clst1 (cluster 1) and clst2 (cluster 2) denote the two clusters obtained in each sample using k-means clustering. P1: UCD46, P2: UCD4, P3: UCD65.

**Figure 3:**
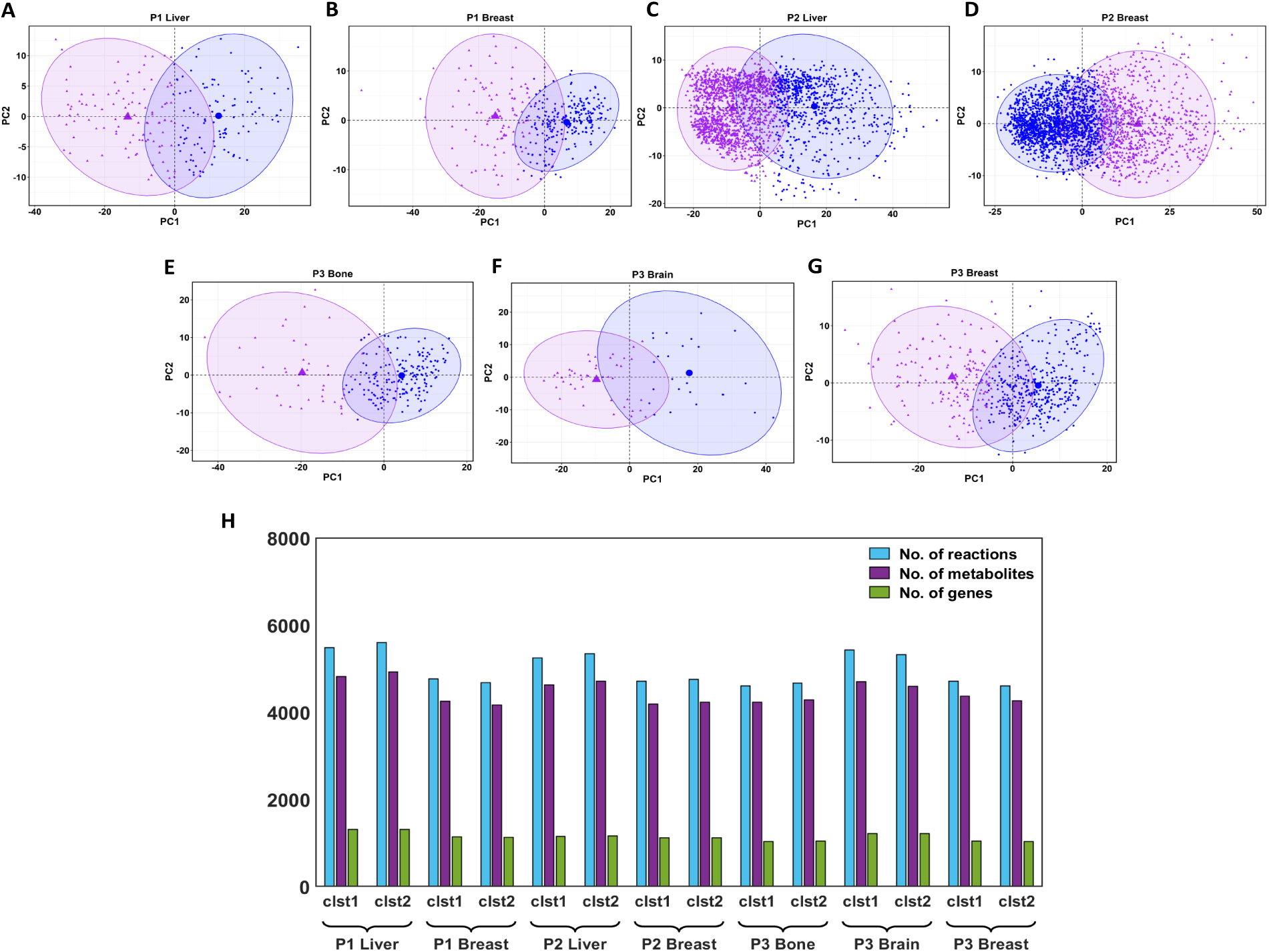
Clustering analysis and metabolic network parameters: (A-G). The figure shows clusters of cells obtained after k-means clustering in each tissue sample. The optimal number of clusters was first determined, and clustering was then performed, revealing distinct groups of cellular profiles based on their feature similarities. (H). The grouped bar plot shows the number of reactions, metabolites, and genes for each cluster-and tissue-specific metabolic models. Here, clst1 and clst2 denote the two clusters obtained in each sample using k-means clustering. P1: UCD46, P2: UCD4, P3: UCD65.

### 2.3 Comparing metabolic profile between primary breast tumor with and without metastatic potential

Next, we wanted to compare the metabolic profiles of the primary breast tumor without metastatic potential and the primary breast tumor with metastatic potential. For this, we chose the GSE235168 dataset, which contains single-cell RNA-seq data from 25 breast cancer patients [30, 31]. In this dataset, two platforms GPL20301 and GPL30173, were used. Among these, the platform GPL20301 was common to the previous dataset, GSE131007, chosen in our study. Therefore, to remove platform-specific biases, we selected patient samples whose single-cell RNA sequencing was performed using the GPL20301 platform. The data underwent filtration and normalization steps, and it was observed that the patients with similar receptor status (i.e., ER*^+^* and Triple-negative breast cancer (TNBC)) were clustered together. Among patients of each subtype, those who clustered relatively more than other patients of that subtype were considered forward. Using k-means, we obtained two optimal clusters within the cells and used the flux value corresponding to each cluster’s centroid as the flux profile for the entire cluster. We next aimed to find the metabolic profile altered in the primary breast tumor with metastatic potential to the liver and bone/brain versus the primary breast tumor without metastatic potential. A cut-off of 1.2 on flux fold change values was applied, and found 393 metabolic reactions were perturbed when comparing the liver-tropic primary tumor (LTPT) with the non-metastatic primary tumor (NMPT), whereas 440 metabolic reactions were perturbed when comparing the bone/brain-tropic primary tumor (BTPT) with the non-metastatic primary tumor (NMPT) (for methodology workflow see Figure S1). Among these, around 200 reactions were common, suggesting a common axis of breast metastasis irrespective of the metastatic site, however, around 49 and 96 reactions were specific to LTPT and BTPT, respectively (see Figure 4A). Most reactions specific to BTPT were down-regulated. There were a few reactions which were perturbed during both cases, i.e., LTPT vs NMPT and BTPT vs NMPT, however, the nature of their alteration was different. Many reactions which were up-regulated in LTPT vs NMPT, were found to be down-regulated in BTPT vs NMPT and vice-versa (see Figure 4B). We further examined the biological pathways enriched by the reactions perturbed between the primary tumors with and without metastatic potential (see Figure 4C). We observed that pathways, namely Nucleotide metabolism, Pyrimidine metabolism, and Transport reactions, showed enrichment of reactions that were upregulated or down-regulated in LTPT vs NMPT and changed their status in BTPT vs NMPT. Whereas, pathways Eicosanoid metabolism, Folate metabolism, Prostaglandin biosynthesis, and Pyruvate metabolism involved reactions that were up-regulated in NMPT vs LTPT and down-regulated in NMPT vs BTPT. Purine metabolism showed involvement of reactions which were down-regulated in LTPT vs NMPT and up-regulated in BTPT vs NMPT.

**Figure 4:**
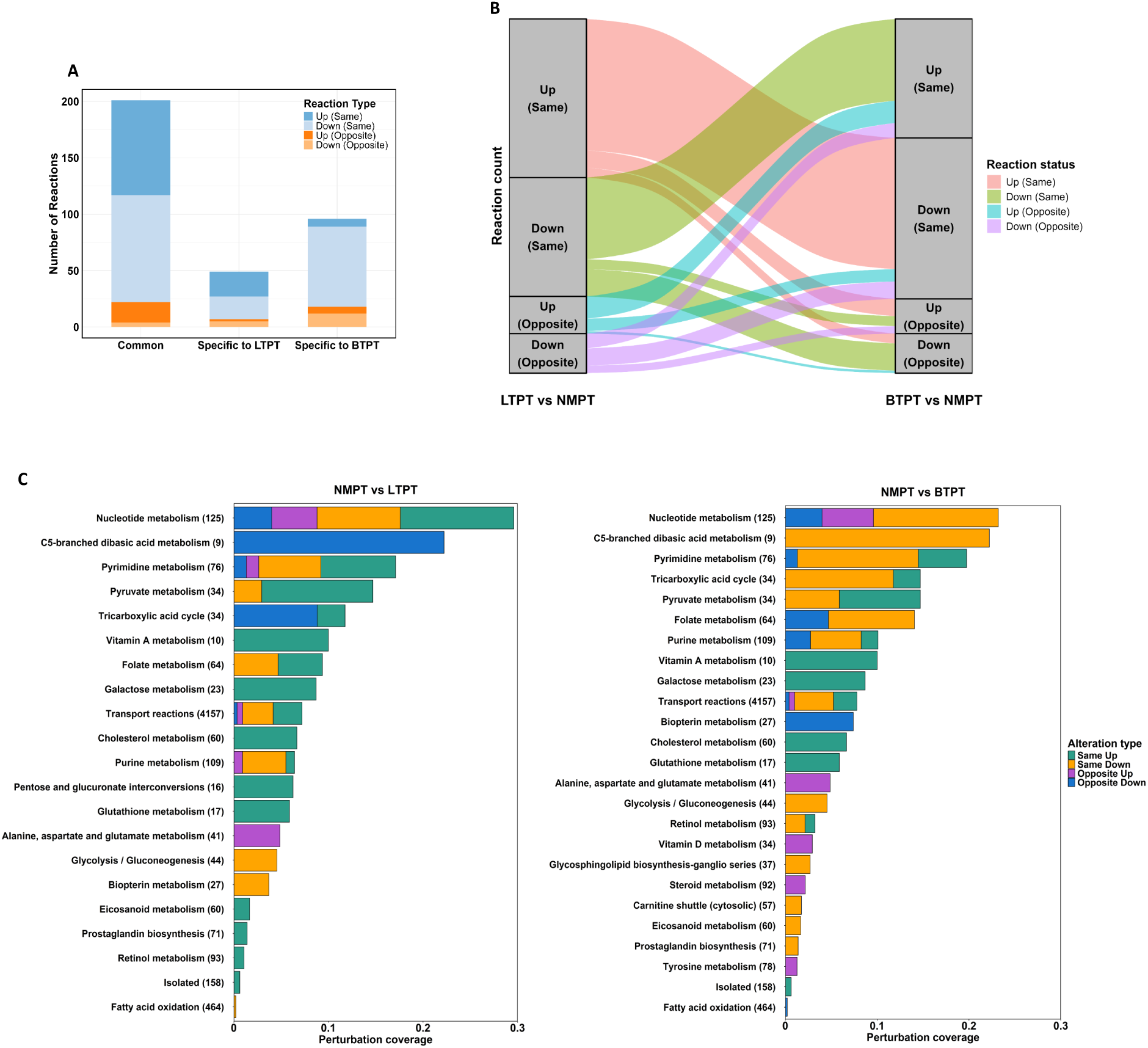
Comparative analysis of primary breast tumors with and without metastatic potential: (A). The stacked bar plot shows the distribution of perturbed reactions comparing the primary breast tumor with and without metastatic potential. (B). Sankey plot displaying the flow of perturbed reactions across the two categories. The left side shows the altered reaction status in LTPT vs NMPT, whereas the right side displays the alteration type in BTPT vs NMPT. The width of each stream is proportional to the number of reactions. (C). Pathway enriched by the reactions perturbed between the primary breast tumor with metastasis to liver & bone, brain vs primary breast tumor without metastasis. The x-axis shows the enriched pathways, and the y-axis shows the ratio of each alteration type in each pathway. The numbers in parentheses along the pathway indicate the total number of reactions in that pathway.

**Figure 5:**
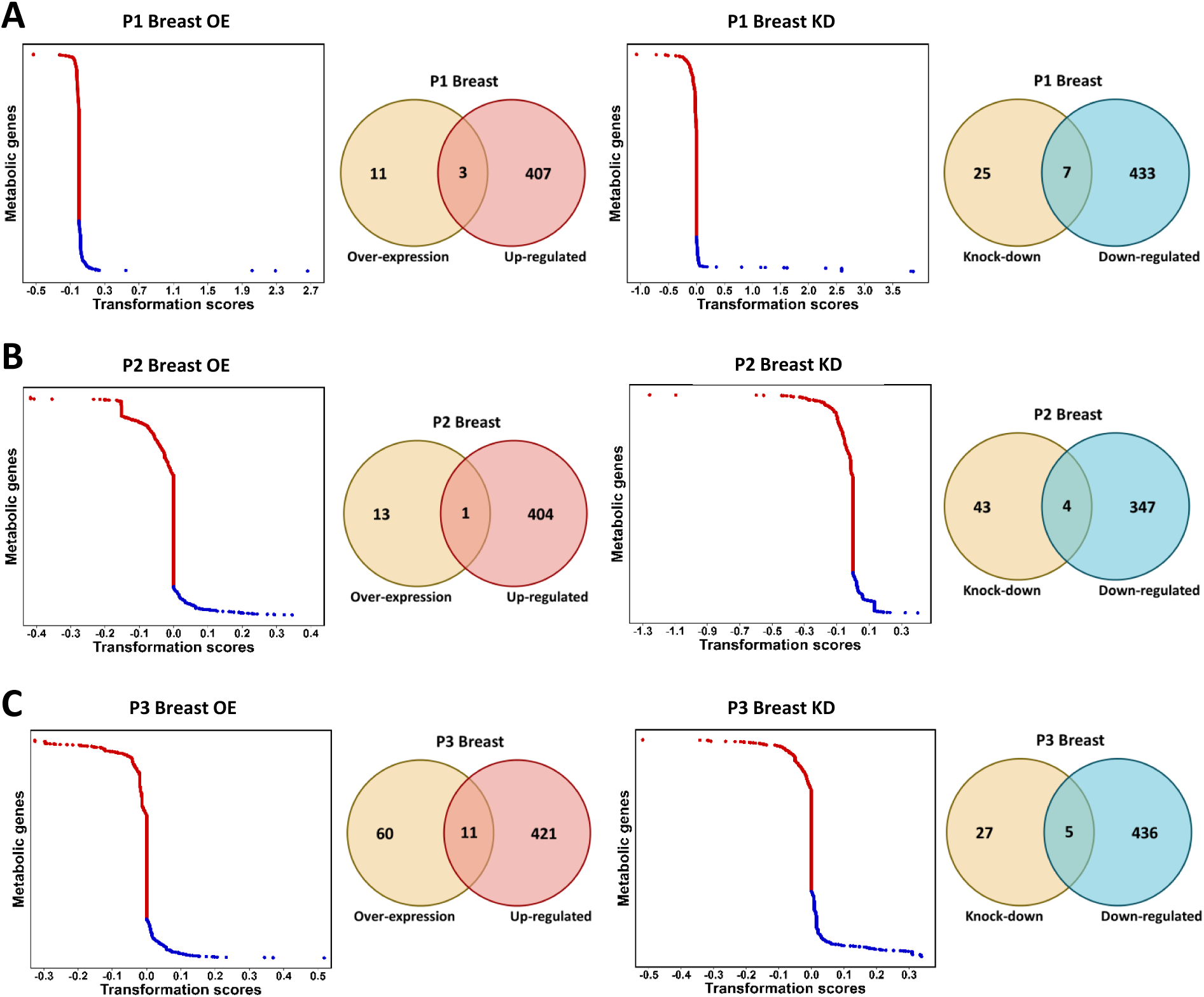
rMTA analysis results: The transformation score of the perturbed metabolic genes obtained by systematically performing a 25%, 50%, 75%, 100% gene over-expression as well as gene knock-down for (A). P1 Breast (B). P2 Breast, (C). P3 Breast. Here, the blue dot represents the genes with transformation score > 0, while the red dot represents the genes with transformation score < 0. The Venn diagram shows the overlap between over-expressed genes and the up-regulated genes in the input data, and between knock-down genes and the down-regulated genes in the input data.

### 2.4 *In silico* exploration of gene over-expression and knock-down to identify metabolic perturbations driving metastatic transformation in primary tumors

Till now, we have studied reaction-level metabolic alterations associated with different breast organotropism. We are now interested in determining metabolic genes whose perturbation (e.g., gene knockdown or overexpression) is predicted to shift breast tumor cell’s metabolism from non-metastatic phenotype (source state) to metastatic phenotype (target state). Therefore, we used robust Metabolic Transformation Analysis (rMTA) to identify metabolic genes responsible for driving the metabolic transformation in primary tumor without metastatic potential, offering mechanistic insight and candidate therapeutic targets. This method involves systematically simulating perturbations (e.g., knock-down/over-expression), comparing flux distributions between two conditions, and finally scoring perturbations by how well they transform the source state’s flux profile towards the target state’s profile. To start with, we first filtered out genes catalysing reactions that are altered in primary breast tumor without metastatic potential compared with primary breast tumor with metastatic potential. Using these genes, we systematically performed a 25%, 50%, 75%, and 100% gene over-expression as well as gene knock-down in the source model which is the model associated with primary breast tumor without metastatic potential, and calculated the transformation score for each over-expression and knock-down (see Figure 10). These transformative scores reflect the extent to which these perturbations altered the metabolic phenotypes. Genes which robustly exhibited positive transformation score for all percentage knock-down were filtered out, representing the genes, which when knocked-down in the primary breast tumor without metastatic ability have the potential to move towards the primary breast tumor with metastatic ability. On the other hand, genes which robustly exhibited a positive transformation score for all percentage over-expression were filtered out, representing the genes that, when over-expressed in the primary breast tumor without metastatic ability have the potential to move towards the primary breast tumor with metastatic ability. The above exercise was done for primary tumor without metastatic potential vs primary tumor with metastatic potential to liver as well as bone and brain across different clusters.

We also checked the overlap between genes identified through rMTA analysis and those differentially expressed between primary breast tumors with and without metastatic potential (see Figure 10). These overlapping genes were further analyzed to explore their biological significance across different breast cancer organotropism revealing distinct patterns of gene-pathway associations (see Table 2). In the P1 Breast KD Down condition, key metabolic genes such as IDH3G, IDH1, ACO2, and IDH3B were associated with bile acid biosynthesis, nicotinate and nicotinamide metabolism, and the tricarboxylic acid cycle, whereas multiple solute carrier (SLC) family members, including SLC3A2, SLC25A1, SLC25A17, and others, were linked specifically to transport reactions. Similar patterns were observed in the P2 Breast KD Down group, highlighting conserved metabolic regulation during breast organotropism to the liver. In the P3 Breast KD Down condition, genes such as GUSB, SORD, and GCLM showed enrichment in degradation pathways, sugar metabolism, and glutathione metabolism, alongside a broad set of nuclear pore complex-and SLC-mediated transport reactions.

**Table 2:**
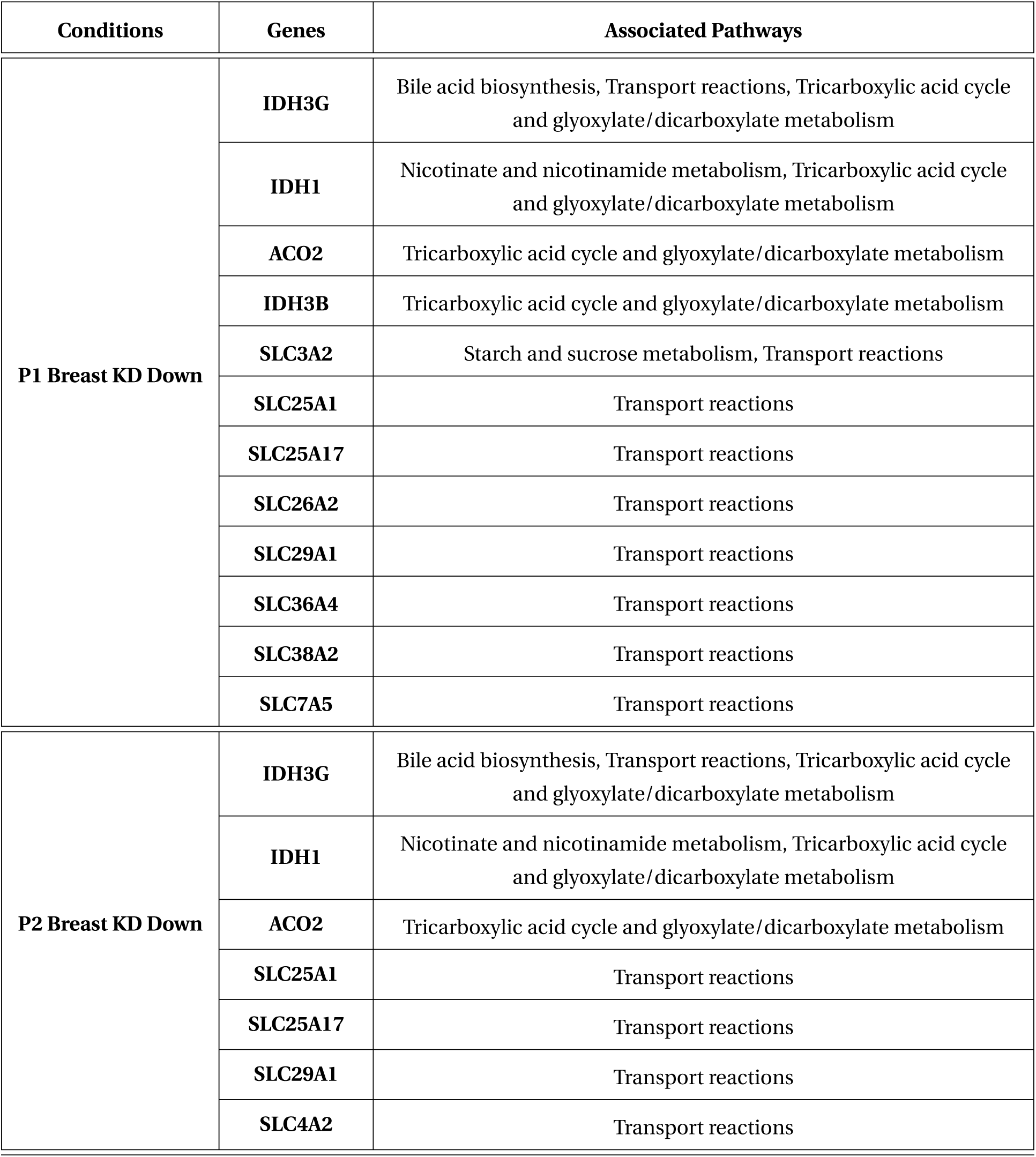
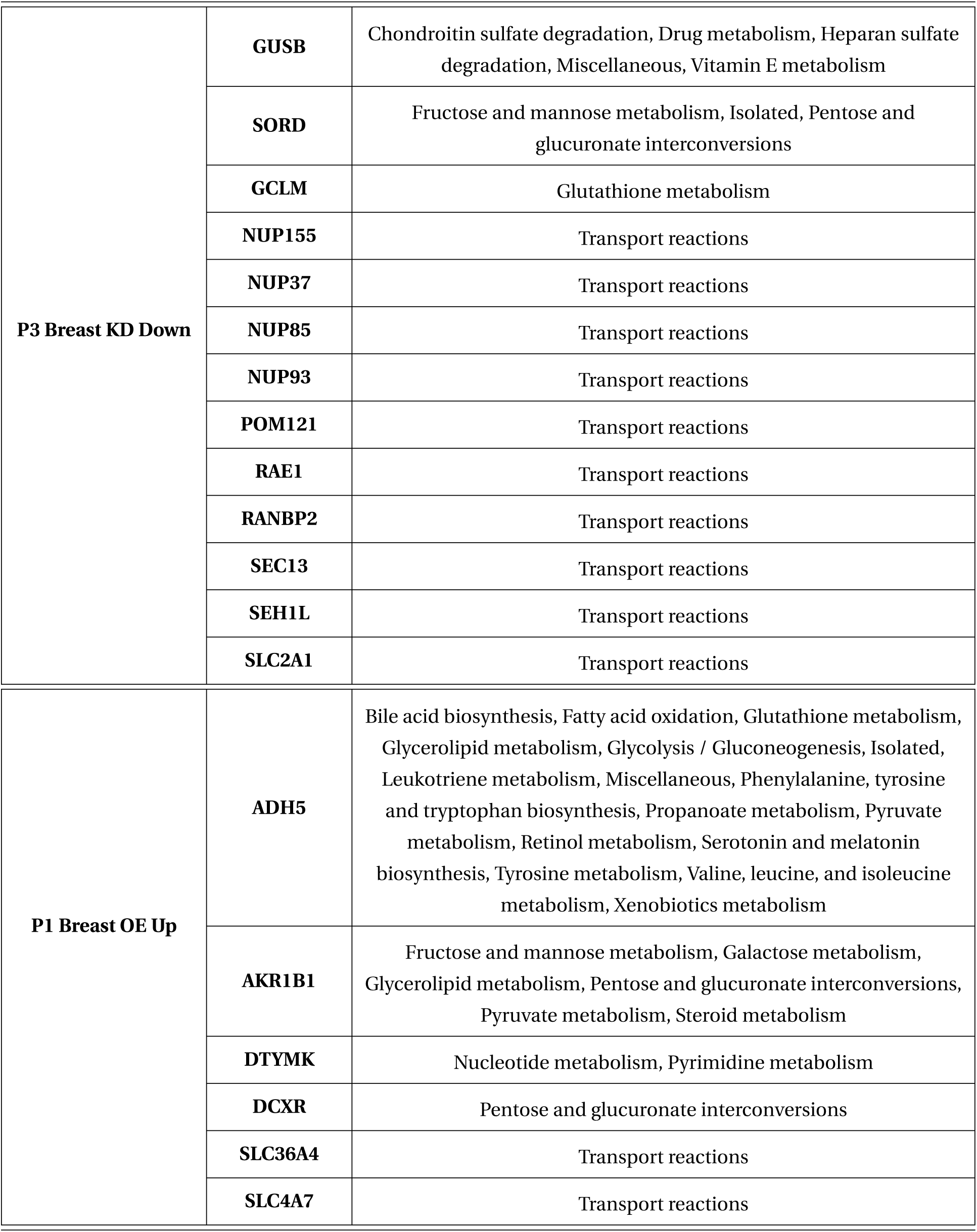
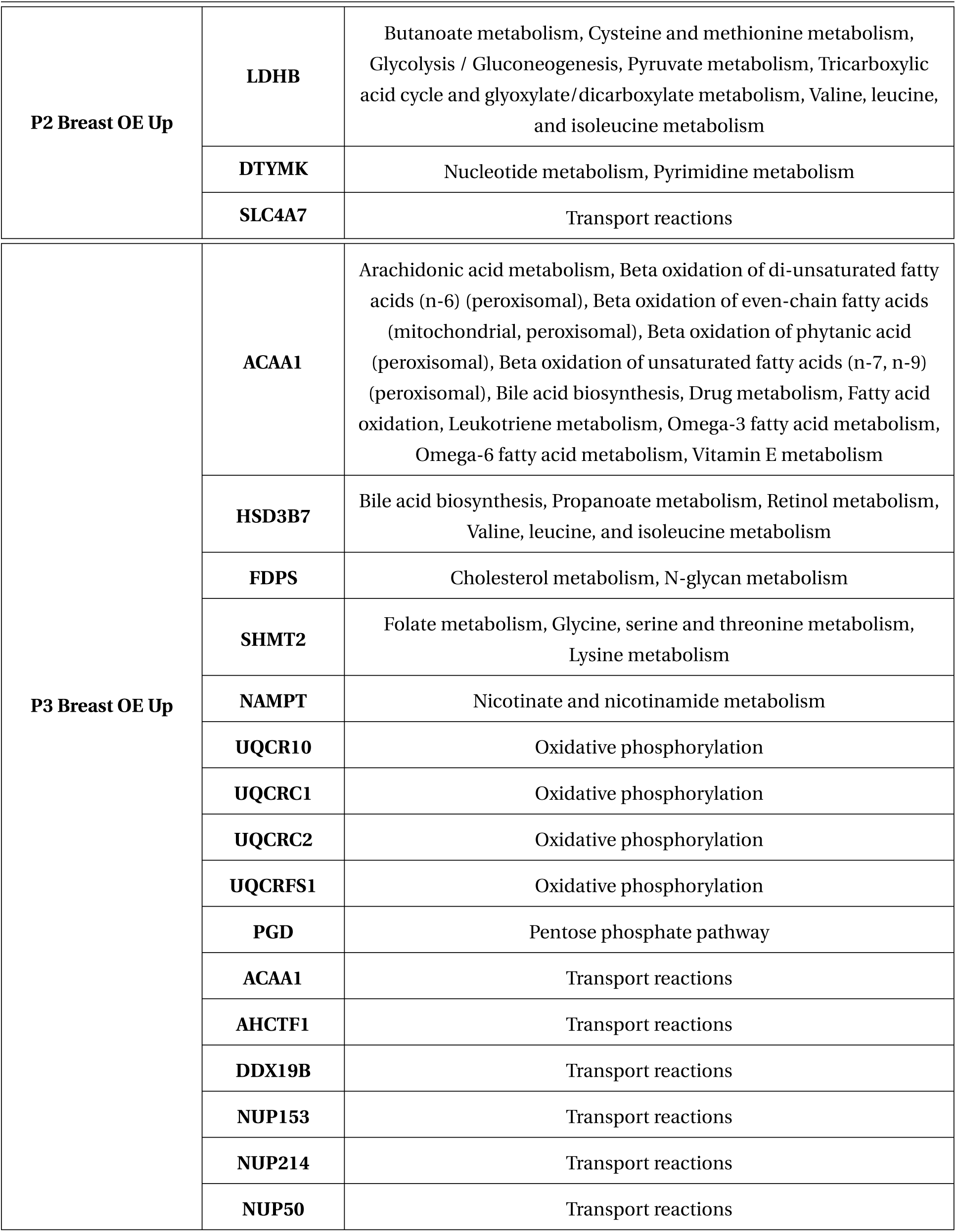
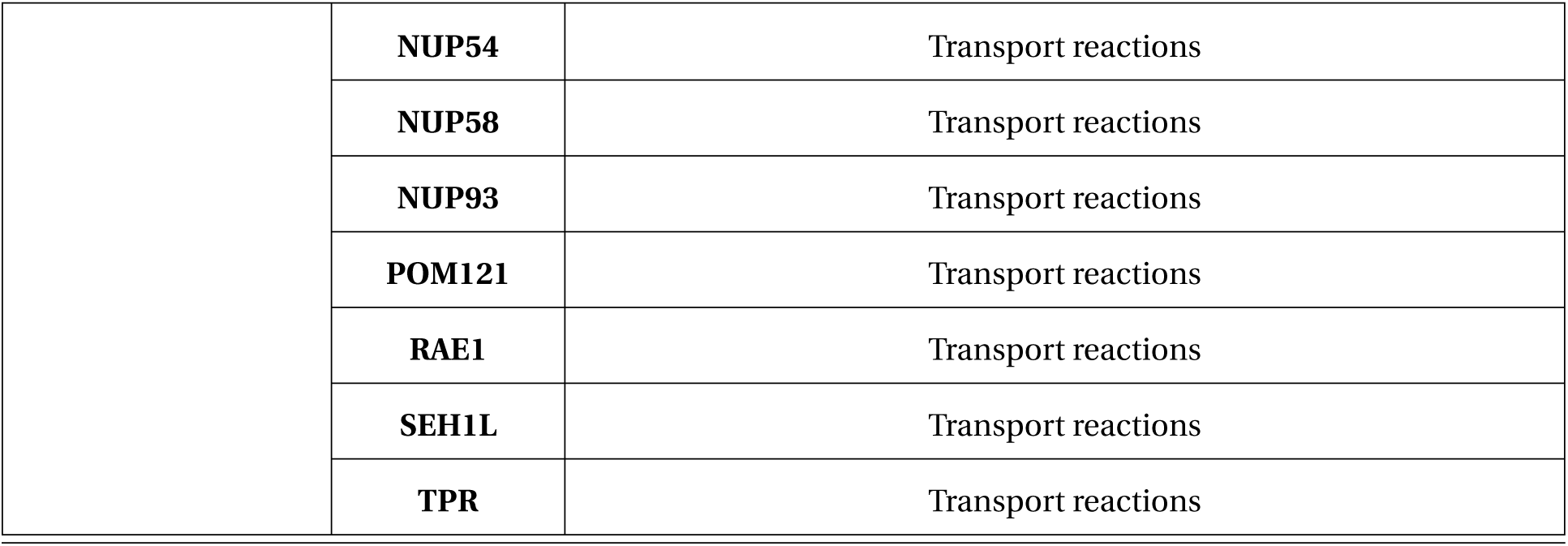
Genes and their associated pathways across breast cancer perturbation conditions: This table lists the genes identified in rMTA analysis for different breast cancer perturbation conditions, including knock-down (KD) and over-expression (OE). For each gene, the associated metabolic or transport pathways are summarized.

Overexpression (OE) perturbations (P1–P3) further revealed extensive up-regulation of metabolic and transport pathways. Notably, ADH5 and AKR1B1 were implicated in diverse biosynthetic and catabolic pathways, including bile acid, fatty acid, glycolysis, and amino acid metabolism, whereas SLC36A4 and SLC4A7 were selectively associated with transport reactions. In P3 Breast OE Up, the analysis uncovered the coordinated involvement of oxidative phosphorylation genes (UQCR10, UQCRC1, UQCRC2, UQCRFS1) and multiple transport genes (ACAA1, AHCTF1, DDX19B, NUP153–NUP93, POM121, RAE1, SEH1L, TPR), emphasizing a potential link between energy metabolism and metabolite transport. Together, these rMTA results identify key metabolic and transport genes whose perturbation may contribute to organ-specific metabolic adaptations in breast cancer.

### 2.5 Validating rMTA targets using CRISPR gene effect data

To further validate our predictions, we integrated results from rMTA analysis with CRISPR-based gene-effect data from the Cancer Dependency Map [32], focusing specifically on breast cancer cell lines. The CRISPR-based gene effect data (CERES scores) measure how a gene knockout impacts cell survival, with a negative score indicating genes essential for survival, meaning their knockout causes cell death, while a positive score represents knocking out the gene increases cell proliferation, suggesting the presence of that gene acts as a brake on growth. Genes with a score around 0 indicate that their knockdown has no effect on proliferation. We compared the experimentally derived gene effect scores with our filtered rMTA-derived perturbations (knockdown and overexpression) that exhibited positive rMTA scores. Specifically, genes showing positive rMTA scores upon knockdown (KD) in the NMPT stage were intersected with genes having positive scores from the CRISPR gene effect dataset of the Cancer Dependency Map. This was based on the rationale that rMTA predicts disease progression upon gene knockdown, and a positive score similarly indicates that gene knockout does not impair, and may even enhance, cell proliferation, thereby supporting disease progression. Using this approach, we identified approximately 11 genes in the case of breast organotropism to liver, and 5 genes in the case of breast organotropism to bone and brain (see Figure 6). The median CRISPR gene effect score for rMTA KD genes was positive for all genes (see Figures 7A,B), and we considered these genes to have a positive gene effect score. Similarly, the median CRISPR gene effect score of rMTA OE genes was found to be negative for all genes (see Figure 7C,D), we considered these as genes with a negative gene effect score. Similarly, genes with positive rMTA scores upon overexpression (OE) in the NMPT stage were intersected with genes exhibiting negative scores. This was motivated by the observation that rMTA predicts disease progression upon gene overexpression, while a negative score indicates that gene knockout reduces cell proliferation, implying that the gene is essential for cancer cell survival. This overlap highlights genes that are both metabolically influential and experimentally essential. Using this criterion, we identified approximately 11 genes in breast organotropism to the liver and 63 genes in breast organotropism to the bone and brain (see Figure 6). Overall, by integrating rMTA predictions with CRISPR-derived gene essentiality, we prioritized metabolically relevant genes that are also experimentally validated to impact cancer cell growth, thereby strengthening the biological relevance and robustness of our findings.

**Figure 6:**
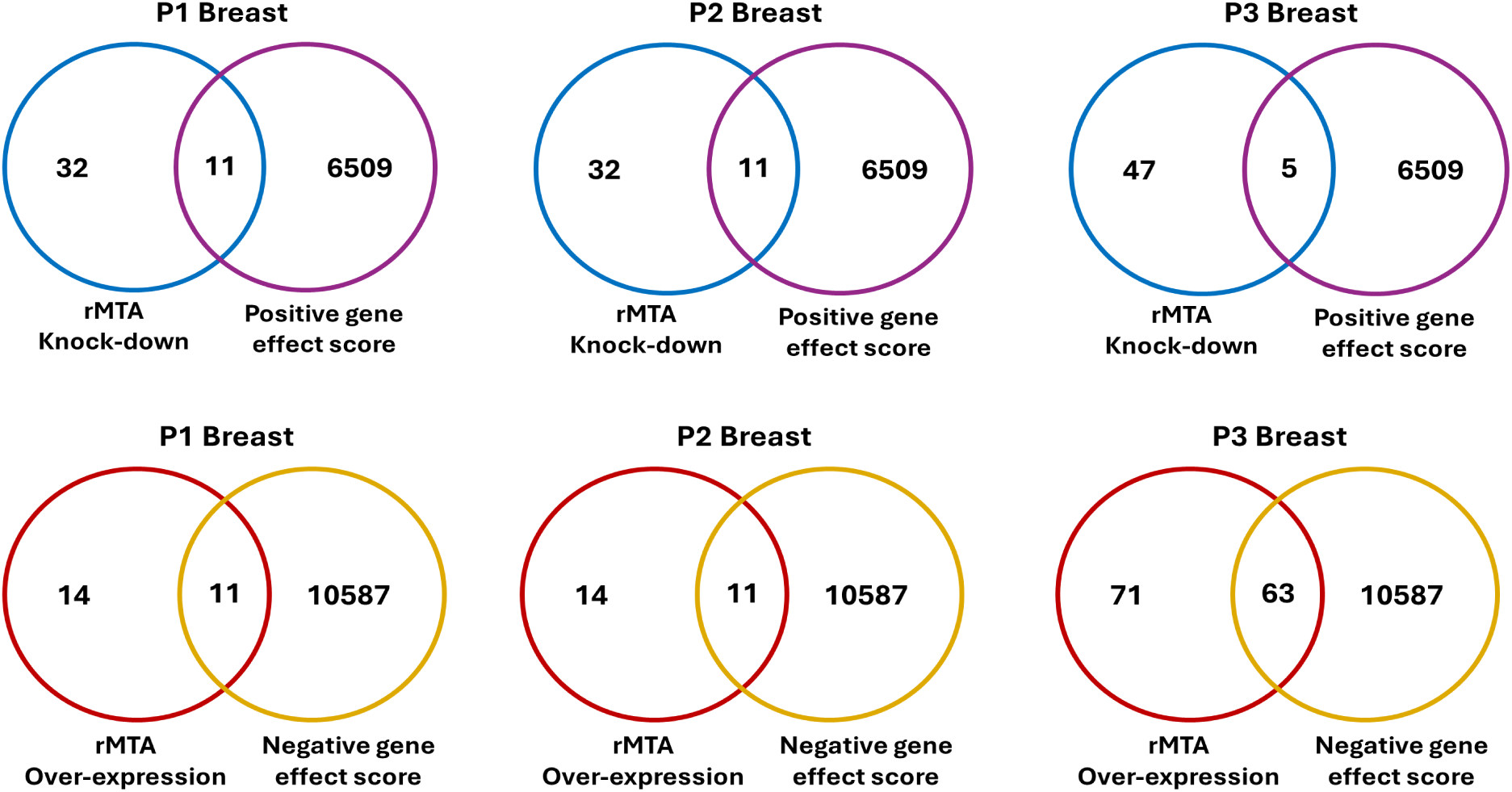
Integration of rMTA-derived perturbations with CRISPR gene effect data: Venn diagrams illustrating the overlap between rMTA-predicted gene perturbations and CRISPR-derived gene effect scores across three breast cancer progression stages (P1, P2, and P3). The top panel represents the intersection of genes with positive rMTA scores upon knockdown (KD) and genes with positive gene effect scores, indicating non-essential genes whose inhibition does not impair cell proliferation and may contribute to disease progression. The bottom panel shows the overlap between genes with positive rMTA scores upon overexpression (OE) and genes with negative gene-effect scores, representing essential genes whose knockouts reduce cell viability, thereby highlighting potential therapeutic targets.

**Figure 7:**
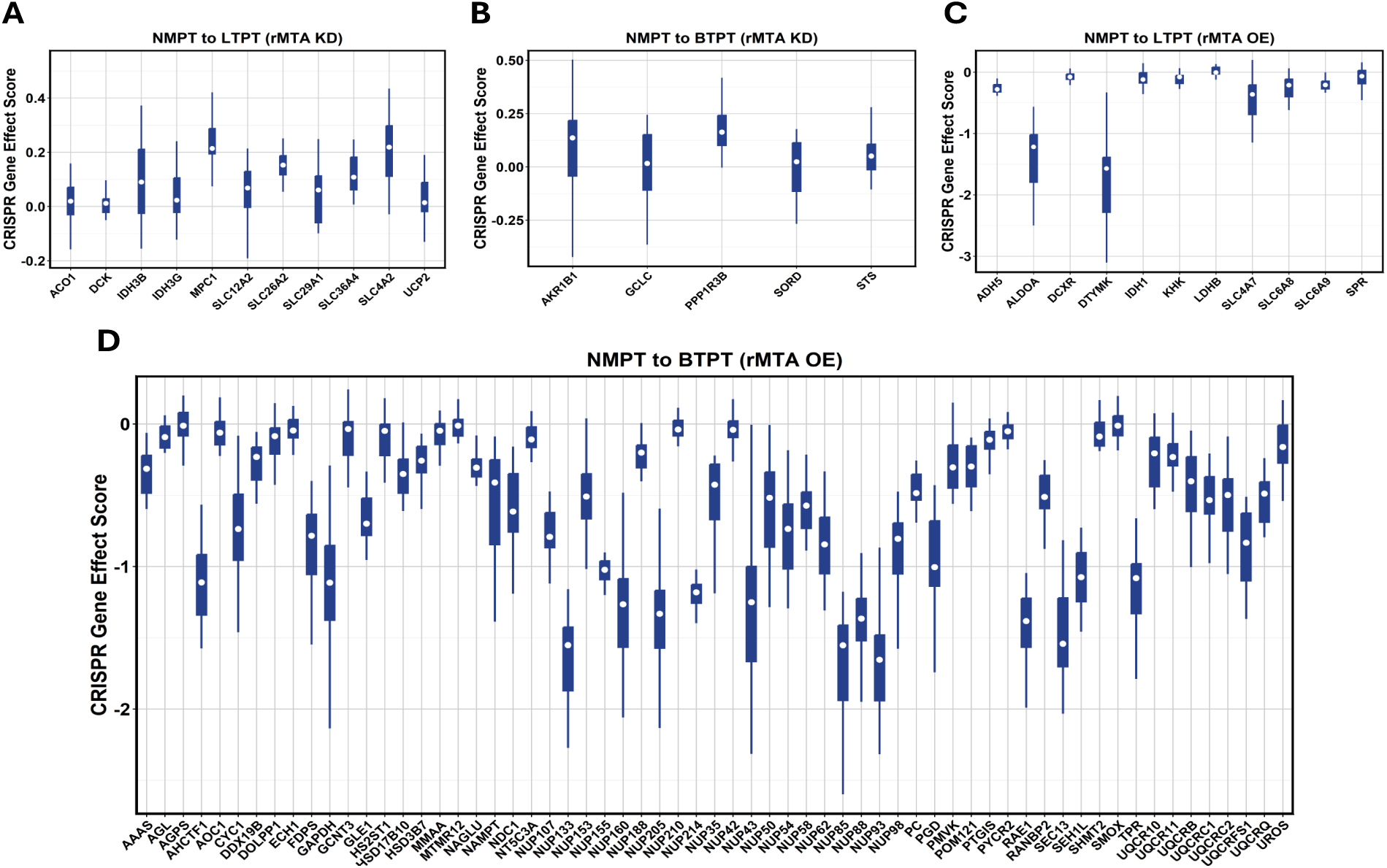
CRISPR gene effect score distribution across rMTA-associated genes: The figure shows the distribution of CRISPR gene effect scores across selected genes associated with rMTA knockdown in (A). NMPT to LTPT, (B). NMPT to BTPT, and rMTA overexpression in (C). NMPT to LTPT, (D). NMPT to BTPT. The boxplots indicate the interquartile range, the white dot denotes the median, and individual data points are overlaid to display the underlying distribution.

### 2.6 Metabolic profiling translates organotropism to the metabolic perturbations

We next aimed to determine which reactions were perturbed during different breast organotropism states. A cut-off of 1.2 was applied to the fold-change values of reactions in the primary breast with metastatic potential and in metastatic organ models to identify reactions perturbed during disease progression for each organotropism (see Figure S2). The number of reactions perturbed during disease transition from primary tumor to each metastatic tumor is shown in Figures 8A-C. Around 100 reactions were commonly perturbed (see Figure 8D), among which up-regulated reactions have a higher share. while others were altered, specific to each organ metastasis. The number of reactions specific to liver metastasis was more than that of bone and brain metastasis. We observed reactions common to all metastases; however, the reaction status varied across them. The majority of the perturbed reactions up-regulated in any two of the metastases shifted their status to down-regulated in the other metastasis (see Figure 8E). Even though these reactions were common, they display different axes for each metastasis, differentiating metabolic liabilities across them.

**Figure 8:**
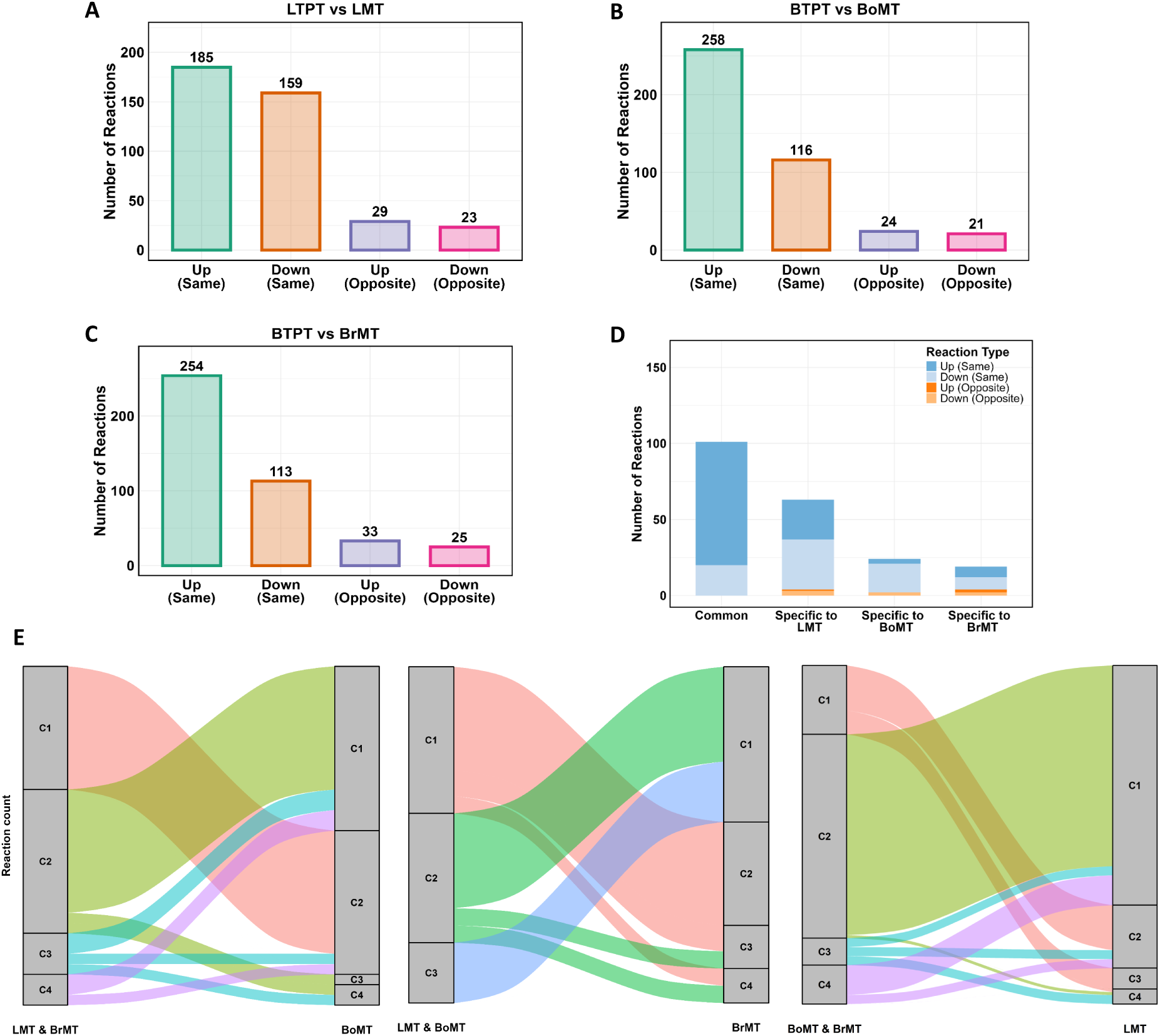
Comparative analysis of primary breast tumors with metastatic potential v/s metastatic secondary organ tumor: The stacked bar plot shows the distribution of perturbed reactions (A). for Liver-tropic primary tumor (LTPT) vs Liver metastatic tumor (LMT). (B). Bone/Brain-tropic primary tumor (BTPT) vs Bone metastatic tumor (BoMT). (C). Bone/Brain-tropic primary tumor (BTPT) vs Brain metastatic tumor (BrMT). (D). The stacked bar plot shows the distribution of perturbed reactions common to all organotropisms and specific to each. (E). Sankey plot displaying the flow of common perturbed reactions with different statuses across the two categories. The left side shows the altered reaction status in two categories, whereas the right side displays the alteration type in the third category. The width of each stream is proportional to the number of reactions. Here, C1: up-regulated reactions (same), C2: down-regulated reactions (same), C3: up-regulated reactions (opposite), C4: down-regulated reactions (opposite).

We analyzed pathways enriched by perturbed reactions shifting their perturbation status across different organotropism (see Tables S2-S7; summarized in Table 3). Reactions up-regulated in the liver and bone but down-regulated in the brain were primarily associated with purine metabolism, folate metabolism, and transport reactions, whereas reactions down-regulated in the liver and bone but up-regulated in the brain were enriched in cholesterol metabolism, pyrimidine metabolism, and transport reactions. Reactions up-regulated in the liver and brain but down-regulated in the bone were enriched in Vitamin A metabolism, purine metabolism, retinol metabolism, nucleotide metabolism, and transport reactions, while the opposite pattern (down in liver and brain, up in bone) highlighted enrichment in pyrimidine metabolism, tyrosine metabolism, nucleotide metabolism, and transport reactions. For reactions up-regulated in bone and brain but down-regulated in the liver, key enriched pathways included the tricarboxylic acid cycle, tyrosine metabolism, nucleotide metabolism, and predominantly transport reactions. Conversely, reactions down-regulated in bone and brain but up-regulated in the liver were enriched in pyruvate metabolism, nucleotide metabolism, and transport reactions. Across all comparisons, transport reactions were consistently enriched. Purine metabolism was common to all cases in which reactions were up-regulated in liver organotropism, whereas pyrimidine metabolism was prevalent in the majority of cases in which reactions were down-regulated in liver organotropism. A few pathways were uniquely enriched depending on the organ and the regulation pattern.

**Table 3:**
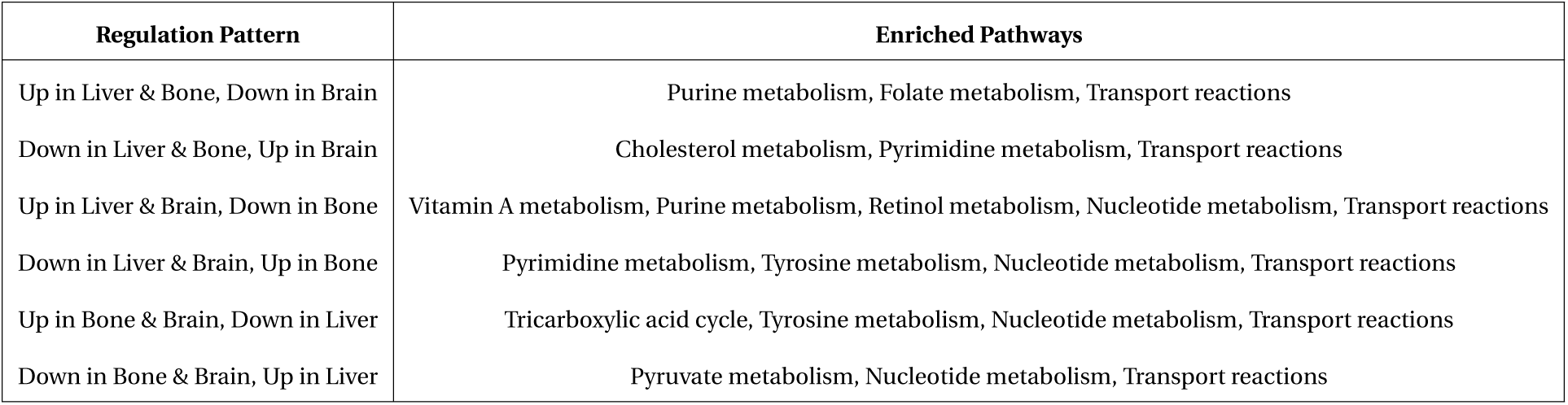
Summary of enriched pathways for perturbed reactions with organ-specific regulation patterns: This table summarizes key metabolic pathways enriched by reactions that exhibit distinct regulation patterns across liver, bone, and brain. Each row indicates a specific up-or down-regulation pattern and the corresponding metabolic pathways enriched in that context.

| Regulation Pattern | Enriched Pathways |
| --- | --- |
| Up in Liver & Bone, Down in Brain | Purine metabolism, Folate metabolism, Transport reactions |
| Down in Liver & Bone, Up in Brain | Cholesterol metabolism, Pyrimidine metabolism, Transport reactions |
| Up in Liver & Brain, Down in Bone | Vitamin A metabolism, Purine metabolism, Retinol metabolism, Nucleotide metabolism, Transport reactions |
| Down in Liver & Brain, Up in Bone | Pyrimidine metabolism, Tyrosine metabolism, Nucleotide metabolism, Transport reactions |
| Up in Bone & Brain, Down in Liver | Tricarboxylic acid cycle, Tyrosine metabolism, Nucleotide metabolism, Transport reactions |
| Down in Bone & Brain, Up in Liver | Pyruvate metabolism, Nucleotide metabolism, Transport reactions |

We further investigated the transport reactions, which were significantly enriched for each case (see Table S8). We observed a considerable number of secretory reactions that facilitate the movement of metabolites from the cytosol to the extracellular space and studied the metabolites and genes associated with these reactions. Such metabolic interactions between tumor cells and their surrounding microenvironment are important in shaping metastatic organotropism, influencing how cancer cells adapt to and colonize distinct organs. In particular, secretory metabolites released by primary breast tumors can remodel the extracellular matrix to facilitate organ-specific colonization. These findings motivated a deeper exploration of the secretory metabolites released by primary breast tumors that potentially drive organ-specific metastatic tropism.

### 2.7 Secretory metabolites during different breast organotropism

To elucidate the metabolic factors underlying breast cancer organotropism, we identified and characterized the secretory metabolites released by primary breast tumors into their microenvironment (see Figure 9A). For this, we analyzed the total flux values of cytosol-to-extracellular reactions associated with secretory metabolites from the primary breast tumor. Comparative flux analysis revealed distinct secretory metabolite profiles associated with primary breast tumors metastasizing to the liver, bone, or brain (see Figure 9B). In the liver-specific context, metabolites such as 5-formyl-THF, 5-methyl-THF, adenosine, cytidine, H_2_O, hydroxide, inosine, THF, thiamin-P, and zinc were specifically secreted. In contrast, L-carnitine and malonate were the only metabolites uniquely secreted in bone and brain organotropism. Among the metabolites commonly secreted across all organotropisms, a high percentage of secretory reactions associated with acetate, *α*-ketoglutarate (AKG), formate, retinoate, butyrate, and pyruvate were specifically perturbed between liver-tropic primary tumors (LTPT) and liver metastatic tumors (LMT), indicating a distinct metabolic adaptation within the liver-tropic context. In contrast, a subset of secretory reactions linked to citrulline and L-carnitine were specifically perturbed between bone/brain-tropic primary tumors (BTPT) and its corresponding metastatic tumors (BoMT and BrMT). Notably, metabolites such as bicarbonate (HCO*^−^*) and malonate were uniquely associated with reactions perturbed between BTPT and BoMT, reflecting specialized metabolic characteristic of the bone niche. Moreover, several metabolites whose secretory reactions were commonly perturbed across all organotropisms, particularly arginine, histidine, lysine, and serine showed a higher degree of perturbations in liver organotropism compared to other metastatic sites. Together, these findings indicate that liver-tropic tumors predominantly secrete folate and nucleotide associated metabolites to support proliferative adaptation, whereas bone and brain-tropic tumors exhibit selective secretion of carnitine and malonate linked metabolites reflecting energy and redox remodeling within these niches.

**Figure 9:**
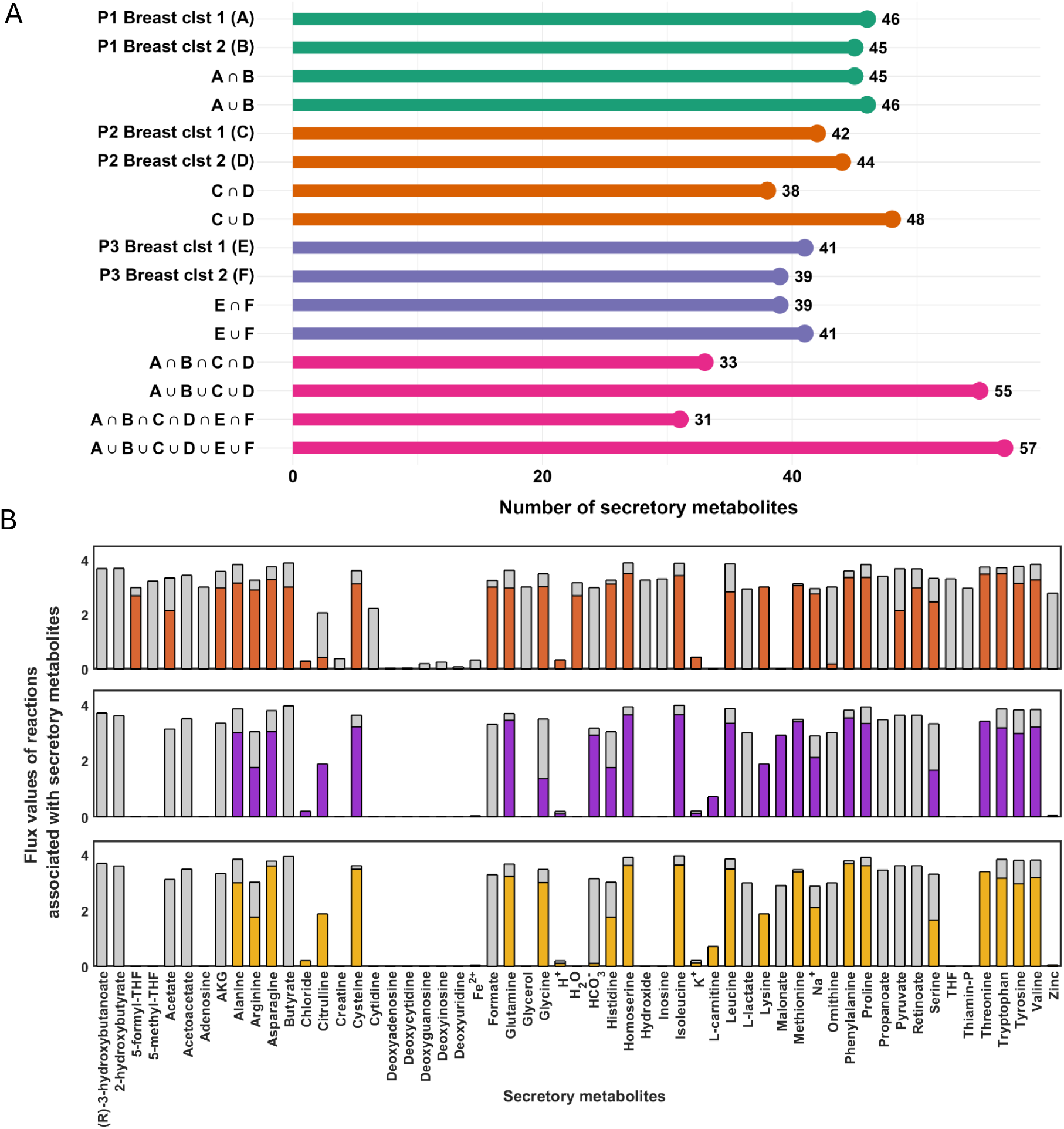
Comparative analysis of flux values of reactions associated with secretory metabolites: (A). The lollipop plot shows the number of secretory metabolites from the cytosol to the extracellular space in each case. (B). The bar graph illustrates secretory metabolite flux values under three different breast organotropism scenarios. Each subplot displays paired bar plots showing total flux (grey bars) and perturbed flux (coloured bars) for each metabolite. In the liver (top panel), perturbed fluxes are highlighted in red, illustrating the effect of specific perturbations on secretory reactions. The bone (middle panel) shows perturbed fluxes in purple, while the brain (bottom panel) uses a yellow color for perturbed reactions. The y-axis in all panels represents the logarithm of the flux values to base 10, incremented by 1 to avoid undefined values at zero. The logarithm to the base 10 transformation was applied to the flux values to scale the flux data, resulting in uniform scaling and allowing direct comparison. The x-axis in all panels represents the individual secretory metabolites.

**Figure 10:**
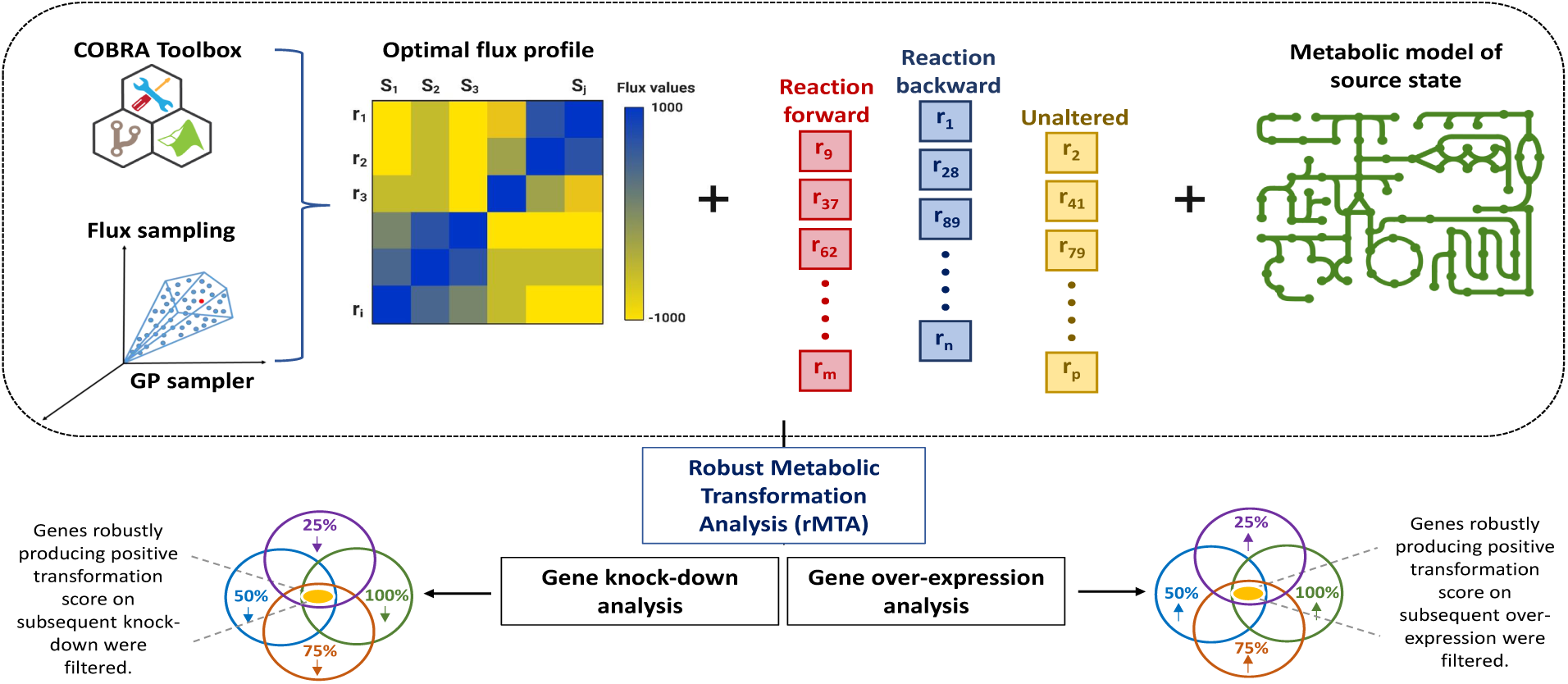
rMTA analysis workflow: The schematic illustrates the workflow of rMTA for identifying key metabolic perturbations associated with metastatic progression. Both *in silico* gene over-expression and knock-down simulations were performed to model transitions from non-metastatic to metastatic primary tumors. Predicted gene targets from rMTA highlight key metabolic differences associated with metastatic potential.

## 3 Materials & Methods

### 3.1 Data acquisition

The public single-cell RNA (scRNA) sequencing data of primary tumors and metastases in mice harbouring breast cancer patient-derived xenografts (PDXs) was retrieved from Gene Expression Omnibus (GEO) with the accession number ‘GSE131007’ [33]. The PDX derived breast cancer cell lines used to study metastasis were UCD4 (invasive ductal carcinoma (IDC), derived from a pleural effusion, ER*^+^*), UCD46 (IDC, derived from primary tumor, ER*^−^*), UCD65 (IDC, lymph node metastasis, ER*^+^*) [34]. These PDX cell lines have been classified into breast cancer subtypes: UCD65 as luminal A, UCD4 as luminal B, and UCD46 as basal-like/triple negative (see Table 4). The data of primary or metastatic tumors which were harvested for scRNA-seq analysis, with liver metastasis collected from UCD4, UCD46 and bone, brain metastases collected from UCD65 are provided in Table 1.

**Table 4:**
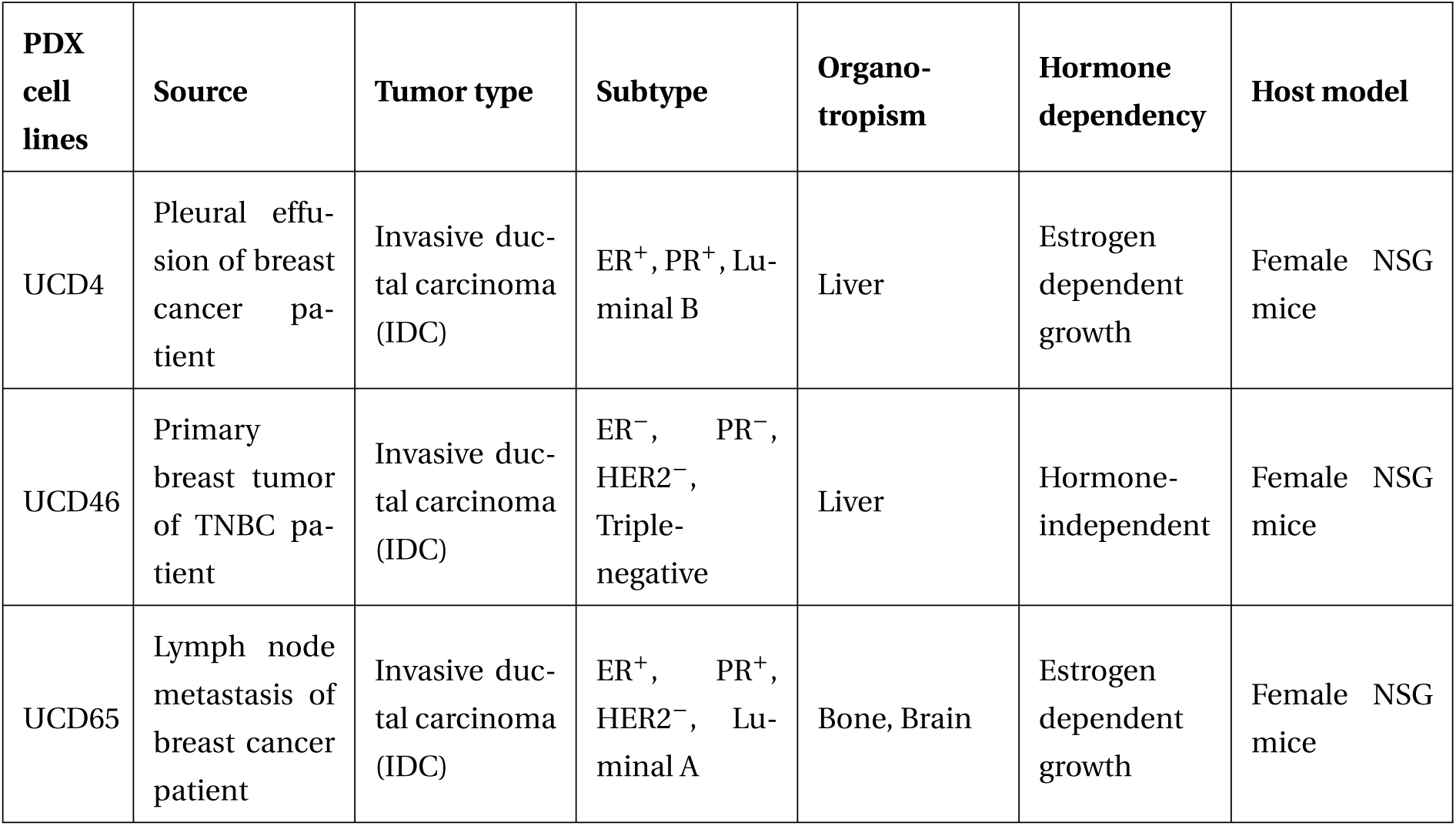
Characteristics of PDX cell lines: Different breast cancer cell lines derived from patient-derived xenografts (PDX) and their general characteristics.

| PDX cell lines | Source | Tumor type | Subtype | Organo-tropism | Hormone dependency | Host model |
| --- | --- | --- | --- | --- | --- | --- |
| UCD4 | Pleural effusion of breast cancer patient | Invasive ductal carcinoma (IDC) | ER<sup>+</sup>, PR<sup>+</sup>, Luminal B | Liver | Estrogen dependent growth | Female NSG mice |
| UCD46 | Primary breast tumor of TNBC patient | Invasive ductal carcinoma (IDC) | ER<sup>-</sup>, PR<sup>-</sup>, HER2<sup>-</sup>, Triple-negative | Liver | Hormone-independent | Female NSG mice |
| UCD65 | Lymph node metastasis of breast cancer patient | Invasive ductal carcinoma (IDC) | ER<sup>+</sup>, PR<sup>+</sup>, HER2<sup>-</sup>, Luminal A | Bone, Brain | Estrogen dependent growth | Female NSG mice |

### 3.2 Quality control and data integration

The public single-cell RNA (scRNA)-seq dataset was downloaded from the Gene Expression Omnibus database, with accession number GSE131007 [33]. First, the quality control was applied to cells: cells were filtered for detected genes (min: 250), mitochondrial gene percent (0–15%), hemoglobin gene percent (0–0.1%) and ribosomal gene percent (min: 1–100%), and total number of molecules detected within a cell (RNA counts > 500), and log_10_Genes PerUMI (the gene numbers of per UMI) less than 0.8 were removed from the dataset [35, 36]. Subsequently, cells were integrated by Canonical Correlation Analysis (CCA) method, using ‘IntegrateData’ function of R package ‘Seurat’ [37]. We next employed the centered log-ratio (CLR) normalization method, which normalizes the read counts using a log transformed normalization factor based on the geometric means of read counts within samples, thereby emphasizing relative abundance and mitigating library size effects [38–41].

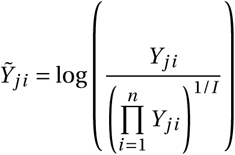

Here, *Y_ji_* represents non-normalized read count for gene or transcript *j* in sample *i*, whereas *Y*^°^*_j i_* represents normalized read counts, *J* represent number of genes and *I* represents the number of samples.

We used Seurat’s FindVariableFeatures() function to identify highly variable genes (HVGs) by modeling the mean–variance relationship of gene expression, selecting genes that capture significant biological heterogeneity for downstream analysis. We applied Seurat’s ScaleData() function to center and scale gene expression values, ensuring that each gene had a mean of zero and unit variance across cells. This step mitigates the influence of genes with large expression ranges and prepares the data for dimensionality reduction and clustering analyses. Then, the UMAP was done for dimension reduction and data visualization. To mark the cell types in the dataset based on gene expression counts, we used the R package ‘celldex,’ which provides a collection of reference expression datasets with curated cell-type labels for use in procedures such as automated annotation of single-cell data [42].

### 3.3 Data pre-processing for model construction

After quality control checks, the scRNA data underwent several steps of data preprocessing. Firstly, epithelial cells that expressed at least 20% of the metabolic genes in each patient’s tissue were filtered out. Then, the metabolic genes expressed in at least 20% of the epithelial cells of each tissue of each patient were filtered. The data were further log-scaled, i.e., log(count + 1). The outliers in the data were identified using the interquartile range (IQR) and corrected using the minimum and maximum values of the corresponding distribution. Next, gene-wise normalisation was performed, i.e., each gene count was divided by the maximum count for that gene across all epithelial cells. We then evaluated the optimal number of clusters in the epithelial cell data for each tissue and patient using the MATLAB function ‘evalclusters’. The cells in each cluster were merged using the centroids of the clusters evaluated through *k*-means clustering.

### 3.4 Reconstruction of tissue-specific metabolic models

The metabolic models specific to each organ and patient considered in the study were extracted using ‘Human1’, which is a generic genome-scale metabolic model for human (Homo sapiens) [43]. The model ‘Human1’ contains 12995 metabolic reactions, 8456 unique metabolites, and 2889 unique metabolic genes distributed over nine subcellular compartments (extracellular, peroxisome, mitochondria, cytosol, lysosome, endoplasmic reticulum, golgi apparatus, nucleus, and inner mitochondria). We generated tissue-specific metabolic models using the Task-driven Integrative Network Inference for Tissues (tINIT) algorithm [28]. The MATLAB function ‘getINIT-Model2’ in the Reconstruction, Analysis and Visualization of Metabolic Networks (RAVEN) Toolbox [44] was utilized to execute the tINIT algorithm, requiring the following inputs:

1. Generic genome-scale metabolic model (Human1)
2. Expression data of metabolic genes in each tissue
3. Gene expression threshold was set as the minimum non-zero value of the metabolic genes in each tissue (above which genes were considered to be ‘expressed’)
4. Structured array of essential metabolic tasks (as returned by function ‘parseTaskList.m’ in the RAVEN Toolbox)

The key steps involved in tINIT algorithm is to assign an expression/score to each reaction, based on evaluating the GPR (Gene-Protein-Reaction) rules using the expression data. The algorithm identifies the core reactions supported by metabolic genes with high expression (above the defined threshold), as well as ensures that the final model is metabolically functional, i.e., capable of performing a set of predefined essential metabolic tasks. To achieve this, the tINIT algorithm builds a Mixed Integer Linear Programming (MILP) problem to maximize the inclusion of reactions supported by high expression and minimizes the inclusion of reactions assigned with low expression score. This results in a pruned, essential metabolic task-performing, expression-consistent context-specific metabolic model.

### 3.5 Mapping of expression data to reactions

The Gene-Protein-Reaction (GPR) rules in ‘Human1’ were utilized to determine the expression data associated with individual metabolic reactions. These rules elucidate the connection between a reaction and the gene product responsible for its catalysis. The ‘GPRparser’ function within the Constrained-Based Reconstruction and Analysis (COBRA) Toolbox [45] transformed these GPR rules into a specific format for subsequent use. The parsed GPR rules, organized in a cell matrix, were then used to integrate the expression data. Utilizing the ‘map-ExpressionToReactions’ function of the COBRA Toolbox facilitated the correlation of expression data with each reaction in the model, resulting in an expression array for each reaction.

### 3.6 Steady state flux profiles evaluation

Constraint-based metabolic models are often underdetermined because they have more reactions than metabolites. Consequently, solutions for such systems encompass a spectrum of possible flux rates rather than a single rate for each reaction. Flux sampling is a common method employed to address genome-scale metabolic models without relying on specific objective functions [46]. This technique produces a series of feasible solutions that adhere to network constraints until the entire solution space is explored. GP sampler, which is a uniform random sampling technique [47], integrated into the COBRA Toolbox was used to account for these feasible solutions. The algorithm iteratively selects a random direction and a random step size in that direction, ensuring the subsequent point remains within the solution space. It efficiently samples the linearly constrained space using a fixed number of points, typically set as 2*number of reactions in the model. The space is defined as:

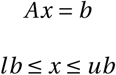

The obtained solution space from the linear constrained problem is represented by:

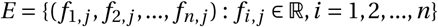

where, *n*= no. of reactions, *j* = fixed no. of points

### 3.7 Optimal flux solution

From the complete sampled solution space, the solution having a maximum mutual information coefficient with the reaction’s expression array was regarded as the optimal flux solution, denoted by:

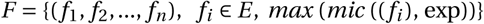

where, *i =* 1, 2, …, *n*, *n*= number of reactions, exp= reaction’s expression array, *mic* ((*f_i_*), exp)= mutual information coefficient (*mic*) between flux values and expression array of metabolic reactions in the queried model. The mutual information between two continuous random variables *X* and *Y* measures the amount of information one variable contains about the other [48]. The MATLAB function ‘*mi*_*cont*_*con*’ was employed to calculate the mutual information, which uses a nearest-neighbours method. The function estimates the joint probability density function (pdf) of *X* and *Y* (*p*^(*x*, *y*)) and marginal pdfs (*p*^(*x*) and *p*^(*y*) of *X* and *Y*, respectively, using k-NN density estimation technique. Using the estimated densities, the mutual information is calculated by:

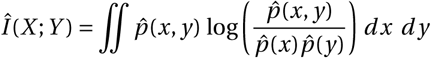

### 3.8 Method to predict metabolic network level perturbation using the gene knockdown and gene overexpression profile

To identify metabolic perturbations capable of transforming the primary metabolic state into the metastatic metabolic state, we performed *in silico* gene knockdown and gene over-expression using the robust Metabolic Transformation Algorithm (rMTA) [49]. The rMTA algorithm extends the original Metabolic Transformation Algorithm (MTA) [50] by evaluating each perturbation under multiple optimization scenarios and integrating the resulting transformation scores into a robust Transformation Score (rTS), thereby providing a more reliable prioritization of candidate metabolic interventions.

The rMTA algorithm requires the following inputs:

1. The reference flux distribution (*V^ref^*) of the source state, estimated using the flux sampling algorithm gpSampler [47]. Among the sampled flux distributions, the flux profile exhibiting the maximum MIC with the corresponding gene expression profile was selected as the reference flux distribution (*V^ref^*).
2. Identification of reactions whose fluxes should change to transform the source metabolic state into the target metabolic state. Based on the reference flux distribution (*V^ref^*), reactions were classified into the forward (*R_F_*), backward (*R_B_*), and unchanged (*R_S_*) reaction sets.

A reaction *r_i_* was assigned to the forward reaction set (*R_F_*) if:

*V_i_^ref^* > 0 and *r_i_* is elevated (up-regulated) or *V_i_^ref^* < 0 and *r_i_* is reduced (down-regulated).

Similarly, a reaction *r_i_* was assigned to the backward reaction set (*R_B_*) if:

*V_i_^ref^* > 0 and *r_i_* is reduced (down-regulated) or *V_i_^ref^* < 0 and *r_i_* is elevated (up-regulated).

The remaining reactions were classified as unchanged (*R_S_*), whose fluxes are expected to remain similar between the source and target metabolic states.

3. Metabolic model of source state.

For each simulated perturbation, the original MTA computes a Transformation Score (TS), which quantifies the ability of the perturbation to transform the source metabolic state towards the target metabolic state. The TS is calculated as

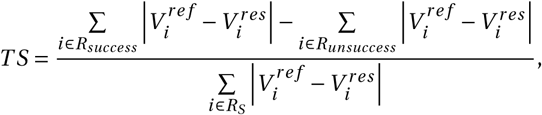

where *V_i_^ref^* and *V_i_^res^* denote the reference and perturbed reaction fluxes, respectively. The altered reactions are classified into *R_success_* and *R_unsuccess_* depending on whether the perturbation changes the reaction flux in the desired or undesired direction. A larger positive TS indicates a greater ability of the perturbation to transform the source metabolic state towards the target metabolic state.

The rMTA algorithm evaluates each perturbation under three complementary optimization scenarios, namely the best-case scenario, yielding the best Transformation Score (*bT S*), the MOMA-based scenario, yielding the MOMA Transformation Score (*mT S*), and the worst-case scenario, yielding the worst Transformation Score (*w T S*). These three scores are subsequently integrated into the robust Transformation Score (*r T S*), which is computed as

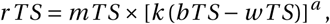

where *k =* 100 is a scaling constant, and

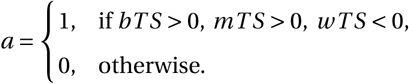

Accordingly, perturbations exhibiting positive *bT S*, positive *mT S*, and negative *w T S* receive an amplified ranking score based on the difference between *bT S* and *w T S*. Otherwise, the exponent becomes zero, reducing the robust Transformation Score to the MOMA Transformation Score (*r T S = mT S*). Consequently, genes with positive rTS values are predicted to consistently shift the source metabolic state towards the target metabolic state and were therefore considered candidate metabolic intervention targets.

For gene knockdown analysis, the upper and lower flux bounds of all reactions associated with a given gene were iteratively reduced by 25%, 50%, 75%, and 100% to simulate different levels of inhibition. Genes exhibiting positive rTS values across all four inhibition levels were retained by intersecting the corresponding gene sets and were considered robust knockdown candidates. Similarly, for gene over-expression analysis, the upper and lower flux bounds of all reactions associated with a given gene were iteratively increased by 25%, 50%, 75%, and 100% to simulate different levels of activation. Genes exhibiting positive rTS values across all four over-expression levels were retained by intersecting the corresponding gene sets and were considered robust over-expression candidates.

The above procedure was independently performed for each source–target cluster transition (*clst1 → clst11*, *clst1 → clst22*, *clst2 → clst11*, and *clst2 → clst22*). Finally, the union of the robust candidate genes identified across all four cluster transitions was used to obtain the final set of metabolic regulators, thereby preserving the metabolic heterogeneity associated with distinct source–target transitions while ensuring robustness of the predicted perturbations across different levels of gene inhibition or activation.

### 3.9 Finding the secretory reactions and metabolites

To identify secretory metabolites, a systematic computational approach was applied using context-specific metabolic models. The optimal flux distribution for each model obtained from context-specific simulations was analyzed to detect active transport reactions allowing metabolite exchange between the cytosolic (‘c’) and extracellular (‘e’) compartments. Reactions were considered secretory if they exhibited a positive flux (above a flux threshold (1 x 10*^−^*^6^) to eliminate numerical noise) in the extracellular direction, indicating net metabolite export. The stoichiometric matrix was used to determine the metabolites participating in each reaction, and only those with positive extracellular coefficients were retained as secretory. The resulting reactions and their associated metabolites were compiled to generate cluster-specific lists of secretory metabolites for each sample. The secretory metabolite results for different clusters corresponding to each organotropism were combined and compared to examine varying secretion patterns of primary breast tumor across different organotropism.

## 4 Discussion

In this study, we investigated the metabolic factors involved in breast cancer organotropism by integrating genome-scale metabolic modeling with single-cell transcriptomics data. This approach allowed us to examine how tumor metabolic programs and organ-specific microenvironments together influence metastatic progression. Our results show that metastatic breast cancer cells undergo different metabolic changes depending on the target organ. Although approximately 200 metabolic reactions were commonly altered across all metastatic trajectories, many metabolic changes were specific to individual organs. These findings suggest the presence of a shared metastatic metabolic program along with organ-specific metabolic adaptations. This combination of common and distinct metabolic features supports the concept of a “core metastatic metabolic axis” shaped by the unique requirements of each metastatic niche. These observations are consistent with the seed-and-soil hypothesis, where intrinsic tumor characteristics (seed) interact with organ-specific microenvironmental conditions (soil) to determine metastatic outcomes [9, 10]. Our findings highlight the importance of context-specific metabolic models for understanding how metabolic adaptations contribute to breast cancer organotropism.

Our comparison between primary tumors with and without metastatic potential revealed that metabolic alterations are already established at the primary site. Pathways such as nucleotide metabolism, pyrimidine metabolism, and transport reactions were consistently perturbed, suggesting that metabolic priming occurs early during tumor progression. This observation is consistent with previous studies demonstrating that metabolic reprogramming is a hallmark of cancer progression and metastasis [14–16]. Interestingly, several pathways exhibited opposite regulation patterns between liver-tropic and bone/brain-tropic tumors, indicating that metabolic divergence toward specific organotropism begins prior to dissemination. This finding underscores the importance of studying primary tumors as predictors of metastatic fate.

The reconstruction of context-specific metabolic models enabled us to capture tissue-and cluster-level heterogeneity in metabolic states. The observed reduction in epithelial cell proportions in metastatic samples, consistent with epithelial–mesenchymal transition (EMT), further emphasizes the interplay between phenotypic plasticity and metabolic adaptation. EMT is well known to facilitate tumor cell invasion, dissemination, and resistance to apoptosis [25]. These transitions likely facilitate both migratory capacity and metabolic flexibility, enabling tumor cells to survive under varying environmental constraints. Through in silico perturbation analysis using rMTA, we identified key metabolic genes whose knockdown or overexpression can drive the transition from a non-metastatic to a metastatic phenotype. Notably, genes associated with the tricarboxylic acid (TCA) cycle (e.g., IDH1, IDH3G, ACO2), redox balance (GCLM), and transport processes (SLC family members) emerged as critical regulators. The prominence of transport-related genes suggests that metabolite exchange between tumor cells and their microenvironment is a central component of metastatic adaptation. This is in line with previous findings emphasizing the importance of metabolic interactions between tumor cells and their niche [26, 51]. Furthermore, the involvement of oxidative phosphorylation genes in overexpression conditions indicates that energy metabolism plays a crucial role in supporting metastatic growth, particularly in energetically demanding environments such as the brain [26].

Integration of rMTA predictions with CRISPR-based gene essentiality data strengthened the biological relevance of our findings. The identification of genes that are both metabolically transformative and experimentally essential highlights potential therapeutic targets. Importantly, differences in the number and type of prioritized genes between liver and bone/brain organotropism suggest that therapeutic strategies may need to be tailored according to metastatic site, consistent with the known heterogeneity of metastatic breast cancer [11–13]. Our analysis of metabolic perturbations during the transition from primary tumors to metastatic sites further revealed that while some reactions are commonly altered, their directionality often differs across organs. This indicates that similar metabolic pathways can be repurposed in distinct ways depending on the target niche. For example, purine metabolism was predominantly associated with liver metastasis, whereas pyrimidine metabolism was more frequently linked to down-regulated processes in the liver context. Such pathway-level distinctions may reflect differences in proliferative demands, nutrient availability, and microenvironmental constraints across organs [20–24, 26].

One of the most striking findings of this study is the identification of organ-specific secretory metabolite profiles. Liver-tropic tumors preferentially secreted folate-and nucleotide-related metabolites, potentially supporting rapid proliferation and biosynthesis in the liver microenvironment. In contrast, bone and brain-tropic tumors exhibited selective secretion of metabolites such as L-carnitine and malonate, which are associated with energy metabolism and redox balance. These secreted metabolites may play a role in pre-metastatic niche formation and in modulating the local microenvironment to favor tumor colonization, as suggested in earlier studies on metastatic niche conditioning [8, 51]. The consistent enrichment of transport reactions across all analyses further supports the importance of metabolic exchange in organotropism.

Several limitations of this study should be acknowledged. First, the analysis was performed using a limited number of patient-derived xenograft samples, which may not fully capture the diversity of breast cancer observed across patient populations. Second, genome-scale metabolic models rely on steady-state assumptions and therefore cannot completely represent the dynamic nature of metabolic regulation in vivo [46]. Third, although our computational analyses identified several candidate genes, pathways, and metabolites, experimental validation will be necessary to confirm their functional roles in metastatic progression.

Future studies should include larger patient cohorts and additional metastatic sites to further evaluate the generalizability of these findings. The integration of other omics data, including proteomics, metabolomics, and spatial transcriptomics, may provide a more complete understanding of the mechanisms driving organotropism [43, 52]. Longitudinal studies following the progression from primary tumor formation to metastatic colonization could also help clarify how metabolic programs evolve over time. Finally, experimental validation in both in vitro and in vivo systems will be essential for translating these findings into clinically relevant therapeutic strategies.

Our study provides a systems-level view of how metabolic adaptation contributes to breast cancer organotropism. The findings suggest that metastatic success depends on the ability of tumor cells to balance conserved metabolic requirements with the specific demands of different organ environments. By revealing metabolic features linked to metastatic site preference, this work expands our understanding of the biological processes that drive organ-specific dissemination. These insights may help guide future studies aimed at developing metabolism-based strategies for controlling metastatic breast cancer.

## Author Contributions

G.A.: Conceptualization (equal); Methodology (lead); Data curation (lead); Formal analysis (lead); Writing - original draft (lead).

S.C.: Conceptualization (equal); Supervision (lead); Writing - review & editing (lead).

## Data Availability Statement

No new data were generated or analysed in support of this research.

## References

[1] Siegel, R.L., Miller, K.D., Wagle, N.S. and Jemal, A., 2023. Cancer statistics, 2023. Ca Cancer J Clin, 73(1), pp.17–48.

[2] Bray, F., Ferlay, J., Soerjomataram, I., Siegel, R.L., Torre, L.A. and Jemal, A., 2018. Global cancer statistics 2018: GLOBOCAN estimates of incidence and mortality worldwide for 36 cancers in 185 countries. CA: a cancer journal for clinicians, 68(6), pp.394–424.

[3] Lukasiewicz, S., Czeczelewski, M., Forma, A., Baj, J., Sitarz, R. and Stanisławek, A., 2021. Breast cancer—epidemiology, risk factors, classification, prognostic markers, and current treatment strategies—an updated review. Cancers, 13(17), p.4287.

[4] Sedeta, E.T., Jobre, B. and Avezbakiyev, B., 2023. Breast cancer: Global patterns of incidence, mortality, and trends.

[5] Mariotto, A.B., Enewold, L., Zhao, J., Zeruto, C.A. and Yabroff, K.R., 2020. Medical care costs associated with cancer survivorship in the United States. Cancer epidemiology, biomarkers & prevention, 29(7), pp.1304–1312.

[6] Lambert, A.W., Pattabiraman, D.R. and Weinberg, R.A., 2017. Emerging biological principles of metastasis. Cell, 168(4), pp.670–691.

[7] Chaffer, C.L. and Weinberg, R.A., 2011. A perspective on cancer cell metastasis. science, 331(6024), pp.1559–1564.

[8] Obenauf, A.C. and Massagué, J., 2015. Surviving at a distance: organ-specific metastasis. Trends in cancer, 1(1), pp.76–91.

[9] Paget, S., 1889. The distribution of secondary growths in cancer of the breast. The Lancet, 133(3421), pp.571–573.

[10] Lu, X. and Kang, Y., 2007. Organotropism of breast cancer metastasis. Journal of mammary gland biology and neoplasia, 12, pp.153–162.

[11] Kennecke, H., Yerushalmi, R., Woods, R., Cheang, M.C.U., Voduc, D., Speers, C.H., Nielsen, T.O. and Gelmon, K., 2010. Metastatic behavior of breast cancer subtypes. Journal of clinical oncology, 28(20), pp.3271–3277.

[12] Chen, W., Hoffmann, A.D., Liu, H. and Liu, X., 2018. Organotropism: new insights into molecular mechanisms of breast cancer metastasis. NPJ precision oncology, 2(1), p.4.

[13] Wu, Q., Li, J., Zhu, S., Wu, J., Chen, C., Liu, Q., Wei, W., Zhang, Y. and Sun, S., 2017. Breast cancer subtypes predict the preferential site of distant metastases: a SEER based study. Oncotarget, 8(17), p.27990.

[14] Gandhi, N. and Das, G.M., 2019. Metabolic reprogramming in breast cancer and its therapeutic implications. Cells, 8(2), p.89.

[15] Faubert, B., Solmonson, A. and DeBerardinis, R.J., 2020. Metabolic reprogramming and cancer progression. Science, 368(6487), p.eaaw5473.

[16] Wang, L., Zhang, S. and Wang, X., 2021. The metabolic mechanisms of breast cancer metastasis. Frontiers in Oncology, 10, p.602416.

[17] Ogrodzinski, M.P., Bernard, J.J. and Lunt, S.Y., 2017. Deciphering metabolic rewiring in breast cancer sub-types. Translational Research, 189, pp.105–122.

[18] Das, C., Bhattacharya, A., Adhikari, S., Mondal, A., Mondal, P., Adhikary, S., Roy, S., Ramos, K., Yadav, K.K., Tainer, J.A. and Pandita, T.K., 2024. A prismatic view of the epigenetic-metabolic regulatory axis in breast cancer therapy resistance. Oncogene, pp.1–15.

[19] Iqbal, M.A., Siddiqui, S., Smith, K., Singh, P., Kumar, B., Chouaib, S. and Chandrasekaran, S., 2023. Metabolic stratification of human breast tumors reveal subtypes of clinical and therapeutic relevance. Iscience, 26(10).

[20] Roda, N., Gambino, V. and Giorgio, M., 2020. Metabolic constrains rule metastasis progression. Cells, 9(9), p.2081.

[21] Bergers, G. and Fendt, S.M., 2021. The metabolism of cancer cells during metastasis. Nature Reviews Cancer, 21(3), pp.162–180.

[22] Elia, I. and Haigis, M.C., 2021. Metabolites and the tumour microenvironment: from cellular mechanisms to systemic metabolism. Nature metabolism, 3(1), pp.21–32.

[23] Rinaldi, G., Rossi, M. and Fendt, S.M., 2018. Metabolic interactions in cancer: cellular metabolism at the interface between the microenvironment, the cancer cell phenotype and the epigenetic landscape. Wiley Interdisciplinary Reviews: Systems Biology and Medicine, 10(1), p.e1397.

[24] Lorusso, G. and Rüegg, C., 2012, June. New insights into the mechanisms of organ-specific breast cancer metastasis. In Seminars in cancer biology (Vol. 22, No. 3, pp. 226–233). Academic Press.

[25] Kalluri, R. and Weinberg, R.A., 2009. The basics of epithelial-mesenchymal transition. The Journal of clinical investigation, 119(6), pp.1420–1428.

[26] Schild, T., Low, V., Blenis, J. and Gomes, A.P., 2018. Unique metabolic adaptations dictate distal organ-specific metastatic colonization. Cancer cell, 33(3), pp.347–354.

[27] Robinson, J.L., Kocabas, P., Wang, H., Cholley, P.E., Cook, D., Nilsson, A., Anton, M., Ferreira, R., Domenzain, I., Billa, V. and Limeta, A., 2020. An atlas of human metabolism. Science signaling, 13(624), p.eaaz1482.

[28] Agren, R., Mardinoglu, A., Asplund, A., Kampf, C., Uhlen, M. and Nielsen, J., 2014. Identification of anti-cancer drugs for hepatocellular carcinoma through personalized genome-scale metabolic modeling. Molecular systems biology, 10(3), p.721.

[29] Sertbas, M. and Ulgen, K.O., 2025. Exploring Human Brain Metabolism via Genome-Scale Metabolic Modeling with Highlights on Multiple Sclerosis. ACS Chemical Neuroscience, 16(7), pp.1346–1360.

[30] Boyd, D.C., Zboril, E.K., Olex, A.L., Leftwich, T.J., Hairr, N.S., Byers, H.A., Valentine, A.D., Altman, J.E., Alzubi, M.A., Grible, J.M. and Turner, S.A., 2023. Discovering synergistic compounds with BYL-719 in PI3K overactivated basal-like PDXs. Cancers, 15(5), p.1582.

[31] Zboril, E.K., Grible, J.M., Boyd, D.C., Hairr, N.S., Leftwich, T.J., Esquivel, M.F., Duong, A.K., Turner, S.A., Ferreira-Gonzalez, A., Olex, A.L. and Sartorius, C.A., 2023. Stratification of tamoxifen synergistic combinations for the treatment of ER+ breast cancer. Cancers, 15(12), p.3179.

[32] Meyers, R.M., Bryan, J.G., McFarland, J.M., Weir, B.A., Sizemore, A.E., Xu, H., Dharia, N.V., Montgomery, P.G., Cowley, G.S., Pantel, S. and Goodale, A., 2017. Computational correction of copy number effect improves specificity of CRISPR–Cas9 essentiality screens in cancer cells. Nature genetics, 49(12), pp.1779–1784.

[33] Dwyer, A.R., Truong, T.H., Kerkvliet, C.P., Paul, K.V., Kabos, P., Sartorius, C.A. and Lange, C.A., 2021. Insulin receptor substrate-1 (IRS-1) mediates progesterone receptor-driven stemness and endocrine resistance in oestrogen receptor+ breast cancer. British Journal of Cancer, 124(1), pp.217–227.

[34] Finlay-Schultz, J., Jacobsen, B.M., Riley, D., Paul, K.V., Turner, S., Ferreira-Gonzalez, A., Harrell, J.C., Kabos, P. and Sartorius, C.A., 2020. New generation breast cancer cell lines developed from patient-derived xenografts. Breast Cancer Research, 22, pp.1–12.

[35] Guo, J., Han, X., Li, J., Li, Z., Yi, J., Gao, Y., Zhao, X. and Yue, W., 2023. Single-cell transcriptomics in ovarian cancer identify a metastasis-associated cell cluster overexpressed RAB13. Journal of Translational Medicine, 21(1), pp.1–15.

[36] Wang, J., Li, X., Zhang, P., Yang, T., Liu, N., Qin, L., Ma, G., Li, X., Fan, H., Huang, S. and Dang, N., 2022. CHRNA5 Is overexpressed in patients with psoriasis and promotes psoriasis-like inflammation in mouse models. Journal of Investigative Dermatology, 142(11), pp.2978–2987.

[37] Stuart, T., Butler, A., Hoffman, P., Hafemeister, C., Papalexi, E., Mauck, W.M., Hao, Y., Stoeckius, M., Smibert, P. and Satija, R., 2019. Comprehensive integration of single-cell data. Cell, 177(7), pp.1888–1902.

[38] Aitchison, J., 1982. The statistical analysis of compositional data. Journal of the Royal Statistical Society: Series B (Methodological), 44(2), pp.139–160.

[39] Quinn, T.P., Crowley, T.M. and Richardson, M.F., 2018. Benchmarking differential expression analysis tools for RNA-Seq: normalization-based vs. log-ratio transformation-based methods. BMC bioinformatics, 19, pp.1–15.

[40] Martino, C., Morton, J.T., Marotz, C.A., Thompson, L.R., Tripathi, A., Knight, R. and Zengler, K., 2019. A novel sparse compositional technique reveals microbial perturbations. MSystems, 4(1), pp.10–1128.

[41] van Lingen, H.J., Suarez-Diez, M. and Saccenti, E., 2024. Normalization of gene counts affects principal components-based exploratory analysis of RNA-sequencing data. Biochimica et Biophysica Acta (BBA)-Gene Regulatory Mechanisms, 1867(4), p.195058.

[42] Aran, D., Looney, A.P., Liu, L., Wu, E., Fong, V., Hsu, A., Chak, S., Naikawadi, R.P., Wolters, P.J., Abate, A.R. and Butte, A.J., 2019. Reference-based analysis of lung single-cell sequencing reveals a transitional profibrotic macrophage. Nature immunology, 20(2), pp.163–172.

[43] Wang, H., Robinson, J.L., Kocabas, P., Gustafsson, J., Anton, M., Cholley, P.E., Huang, S., Gobom, J., Svensson, T., Uhlen, M. and Zetterberg, H., 2021. Genome-scale metabolic network reconstruction of model animals as a platform for translational research. Proceedings of the National Academy of Sciences, 118(30), p.e2102344118.

[44] Wang, H., Marcišauskas, S., Sánchez, B.J., Domenzain, I., Hermansson, D., Agren, R., Nielsen, J. and Kerkhoven, E.J., 2018. RAVEN 2.0: A versatile toolbox for metabolic network reconstruction and a case study on Streptomyces coelicolor. PLoS computational biology, 14(10), p.e1006541.

[45] Heirendt, L., Arreckx, S., Pfau, T., Mendoza, S.N., Richelle, A., Heinken, A., Haraldsdóttir, H.S., Wachowiak, J., Keating, S.M., Vlasov, V., et al. (2019). Creation and analysis of biochemical constraint-based models using the COBRA Toolbox v.3.0. Nat. Protoc. 14, 639–702. 10.1038/s41596-018-0098-2.

[46] Herrmann, H.A., Dyson, B.C., Vass, L., Johnson, G.N., and Schwartz, J.-M. (2019). Flux sampling is a powerful tool to study metabolism under changing environmental conditions. NPJ Syst Biol Appl 5, 32. 10.1038/s41540-019-0109-0.

[47] Schellenberger, J., and Palsson, B.Ø. (2009). Use of randomized sampling for analysis of metabolic networks. J. Biol. Chem. 284, 5457–5461. 10.1074/jbc.R800048200.

[48] Kraskov, A., Stögbauer, H. and Grassberger, P., 2004. Estimating mutual information. Physical review E, 69(6), p.066138.

[49] Valcarcel, L.V., Torrano, V., Tobalina, L., Carracedo, A. and Planes, F.J., 2019. rMTA: robust metabolic transformation analysis. Bioinformatics, 35(21), pp.4350–4355.

[50] Yizhak, K., Gabay, O., Cohen, H. and Ruppin, E., 2013. Model-based identification of drug targets that revert disrupted metabolism and its application to ageing. Nature communications, 4(1), p.2632.

[51] Celia-Terrassa, T. and Kang, Y., 2018. Metastatic niche functions and therapeutic opportunities. Nature cell biology, 20(8), pp.868–877.

[52] Gustafsson, J., Anton, M., Roshanzamir, F., Jörnsten, R., Kerkhoven, E.J., Robinson, J.L. and Nielsen, J., 2023. Generation and analysis of context-specific genome-scale metabolic models derived from single-cell RNA-Seq data. Proceedings of the National Academy of Sciences, 120(6), p.e2217868120.

