## Supplemental File for "Single-cell informed metabolic modeling reveals organ-specific metabolic adaptations in breast cancer organotropism"

### Supplementary Figures

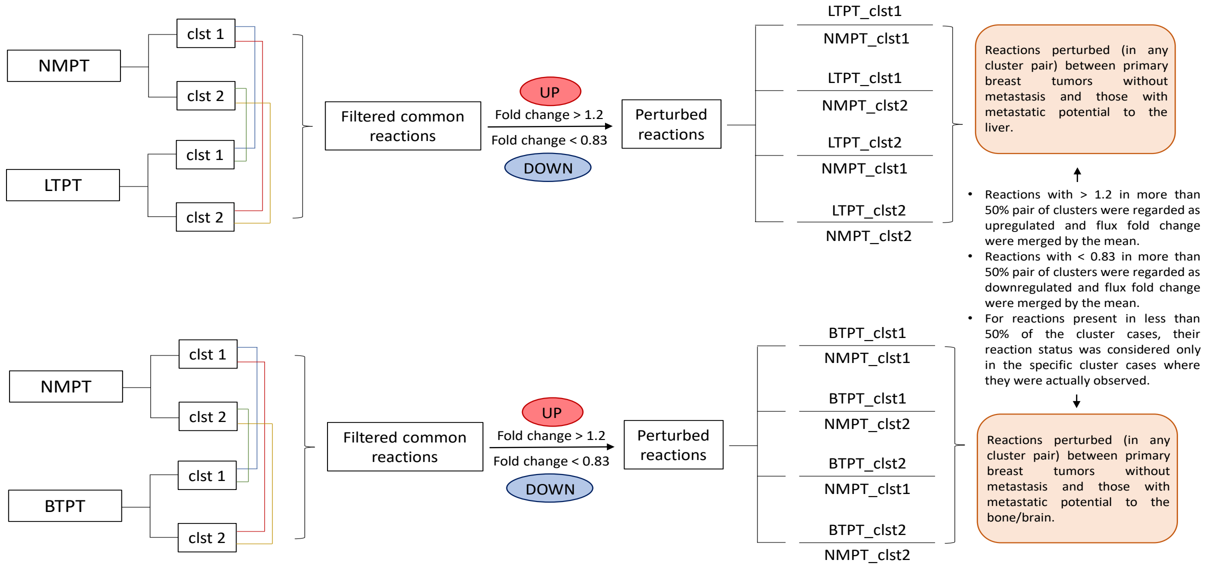


Figure S1: **Comparing primary breast tumors with and without metastatic potential:** The workflow shows the methodology followed to study the perturbations between primary breast tumors with and without metastatic potential.

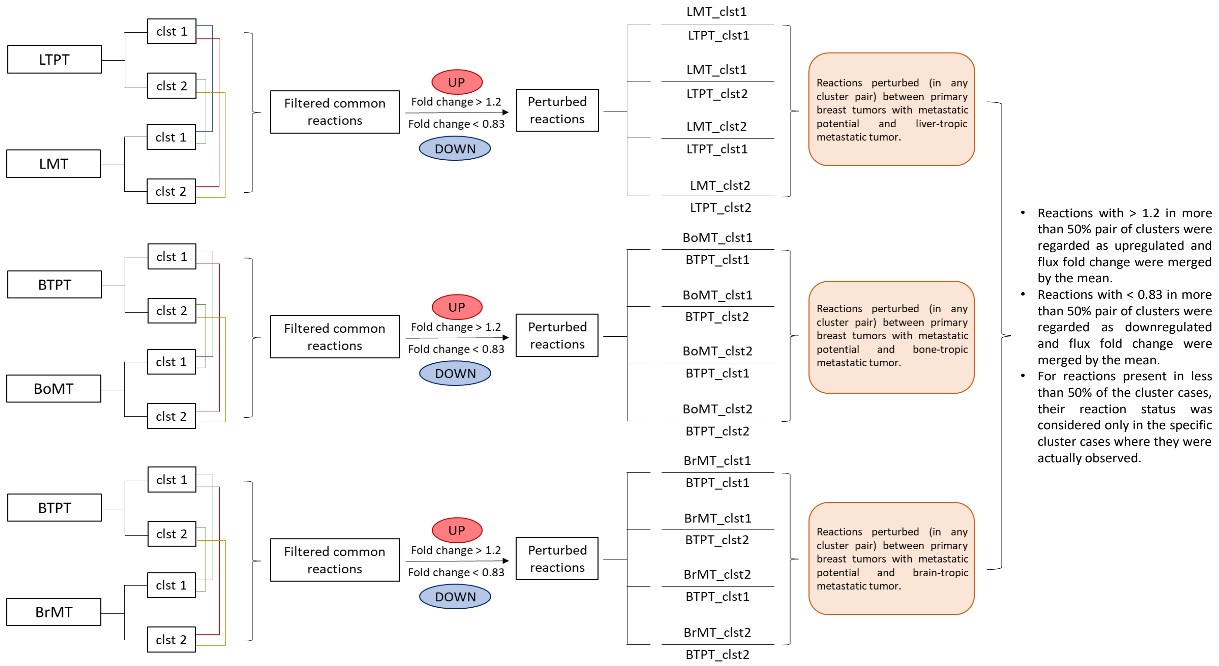


Figure S2: **Primary breast tumor with metastasis versus metastatic secondary organ tumor:** The workflow shows the methodology followed to study the perturbations between primary breast tumors with metastatic potential and their corresponding metastatic tumors.

### Supplementary Tables

Table S1: **Metabolic network parameters for each context-specific model:** The table lists the total number of reactions, associated genes, and metabolites in each model. The metabolic functionality score (out of 57) indicates the completeness of the network in carrying out essential metabolic tasks. Differences in reaction, gene, and metabolite counts reflect subtle context-specific variations in metabolic network architecture across tissues and patients.

| **Context-specific metabolic models** | **No. of reactions** | **No. of genes** | **No. of metabolites** | **Metabolic functionality score (out of 57)** |
|---|---|---|---|---|
| P1 Liver clst 1 | 5480 | 1305 | 4820 | 57 |
| P1 Liver clst 2 | 5599 | 1311 | 4926 | 57 |
| P1 Breast clst 1 | 4765 | 1134 | 4252 | 57 |
| P1 Breast clst 2 | 4684 | 1123 | 4166 | 57 |
| P2 Liver clst 1 | 5250 | 1149 | 4628 | 57 |
| P2 Liver clst 2 | 5338 | 1159 | 4710 | 57 |
| P2 Breast clst 1 | 4706 | 1115 | 4182 | 57 |
| P2 Breast clst 2 | 4757 | 1115 | 4229 | 57 |
| P3 Bone clst 1 | 4609 | 1034 | 4230 | 57 |
| P3 Bone clst 2 | 4671 | 1048 | 4285 | 57 |
| P3 Brain clst 1 | 5426 | 1215 | 4698 | 57 |
| P3 Brain clst 2 | 5326 | 1213 | 4597 | 57 |
| P3 Breast clst 1 | 4714 | 1046 | 4365 | 57 |
| P3 Breast clst 2 | 4608 | 1032 | 4266 | 57 |

Table S2: List of commonly perturbed reactions with same perturbation status in Liver and Bone (i.e., Up-regulated) but different in Brain (i.e., Down-regulated).

| **Reaction ID** | **Liver** | **Bone** | **Brain** | **Pathways** |
|---|---|---|---|---|
| MAR02192 | Same_Up | Same_Up | Same_Down | Miscellaneous |
| MAR02195 | Same_Up | Same_Up | Same_Down | Miscellaneous |
| MAR04333 | Same_Up | Same_Up | Same_Down | Folate metabolism |
| MAR04421 | Same_Up | Same_Up | Same_Down | Purine metabolism |
| MAR05470 | Same_Up | Same_Up | Same_Down | Transport reactions |
| MAR05561 | Same_Up | Same_Up | Same_Down | Transport reactions |
| MAR05607 | Same_Up | Same_Up | Same_Down | Transport reactions |
| MAR05629 | Same_Up | Same_Up | Same_Down | Transport reactions |
| MAR05660 | Same_Up | Same_Up | Same_Down | Transport reactions |
| MAR05673 | Same_Up | Same_Up | Same_Down | Transport reactions |
| MAR20125 | Same_Up | Same_Up | Same_Down | Transport reactions |
| MAR20166 | Same_Up | Same_Up | Same_Down | Transport reactions |

Table S3: List of commonly perturbed reactions with same perturbation status in Liver and Bone (i.e., Down-regulated) but different in Brain (i.e., Up-regulated).

| **Reaction ID** | **Liver** | **Bone** | **Brain** | **Pathways** |
|---|---|---|---|---|
| MAR00443 | Same_Down | Same_Down | Same_Up | Cholesterol metabolism |
| MAR02206 | Same_Down | Same_Down | Same_Up | Cholesterol metabolism |
| MAR02337 | Same_Down | Same_Down | Same_Up | Cholesterol metabolism |
| MAR02339 | Same_Down | Same_Down | Same_Up | Cholesterol metabolism |
| MAR04673 | Same_Down | Same_Down | Same_Up | Pyrimidine metabolism |
| MAR04675 | Same_Down | Same_Down | Same_Up | Pyrimidine metabolism |
| MAR05074 | Same_Down | Same_Down | Same_Up | Transport reactions |
| MAR05487 | Same_Down | Same_Down | Same_Up | Transport reactions |
| MAR05659 | Same_Down | Same_Down | Same_Up | Transport reactions |
| MAR05674 | Same_Down | Same_Down | Same_Up | Transport reactions |
| MAR20147 | Same_Down | Same_Down | Same_Up | Transport reactions |

Table S4: List of commonly perturbed reactions with same perturbation status in Liver and Brain (i.e., Up-regulated) but different in Bone (i.e., Down-regulated).

| **Reaction ID** | **Liver** | **Bone** | **Brain** | **Pathways** |
|---|---|---|---|---|
| MAR01690 | Same_Up | Same_Down | Same_Up | Vitamin A metabolism |
| MAR01917 | Same_Up | Same_Down | Same_Up | Transport reactions |
| MAR04044 | Same_Up | Same_Down | Same_Up | Purine metabolism |
| MAR04863 | Same_Up | Same_Down | Same_Up | Transport reactions |
| MAR05046 | Same_Up | Same_Down | Same_Up | Transport reactions |
| MAR05580 | Same_Up | Same_Down | Same_Up | Transport reactions |
| MAR05625 | Same_Up | Same_Down | Same_Up | Transport reactions |
| MAR06644 | Same_Up | Same_Down | Same_Up | Retinol metabolism |
| MAR07638 | Same_Up | Same_Down | Same_Up | Transport reactions |
| MAR07873 | Same_Up | Same_Down | Same_Up | Nucleotide metabolism |
| MAR20140 | Same_Up | Same_Down | Same_Up | Transport reactions |
| MAR20150 | Same_Up | Same_Down | Same_Up | Transport reactions |

Table S5: List of commonly perturbed reactions with same perturbation status in Liver and Brain (i.e., Down-regulated) but different in Bone (i.e., Up-regulated).

| **Reaction ID** | **Liver** | **Bone** | **Brain** | **Pathways** |
|---|---|---|---|---|
| MAR02414 | Same_Down | Same_Up | Same_Down | Transport reactions |
| MAR04008 | Same_Down | Same_Up | Same_Down | Pyrimidine metabolism |
| MAR04030 | Same_Down | Same_Up | Same_Down | Pyrimidine metabolism |
| MAR04635 | Same_Down | Same_Up | Same_Down | Pyrimidine metabolism |
| MAR04683 | Same_Down | Same_Up | Same_Down | Tyrosine metabolism |
| MAR04686 | Same_Down | Same_Up | Same_Down | Tyrosine metabolism |
| MAR05324 | Same_Down | Same_Up | Same_Down | Transport reactions |
| MAR05645 | Same_Down | Same_Up | Same_Down | Transport reactions |
| MAR05664 | Same_Down | Same_Up | Same_Down | Transport reactions |
| MAR05873 | Same_Down | Same_Up | Same_Down | Transport reactions |
| MAR08450 | Same_Down | Same_Up | Same_Down | Nucleotide metabolism |
| MAR20128 | Same_Down | Same_Up | Same_Down | Transport reactions |

Table S6: List of commonly perturbed reactions with same perturbation status in Bone and Brain (i.e., Up-regulated) but different in Liver (i.e., Down-regulated).

| **Reaction ID** | **Liver** | **Bone** | **Brain** | **Pathways** |
|---|---|---|---|---|
| MAR03958 | Same_Down | Same_Up | Same_Up | Tricarboxylic acid cycle |
| MAR04685 | Same_Down | Same_Up | Same_Up | Tyrosine metabolism |
| MAR04687 | Same_Down | Same_Up | Same_Up | Tyrosine metabolism |
| MAR04852 | Same_Down | Same_Up | Same_Up | Transport reactions |
| MAR05076 | Same_Down | Same_Up | Same_Up | Transport reactions |
| MAR05077 | Same_Down | Same_Up | Same_Up | Transport reactions |
| MAR05084 | Same_Down | Same_Up | Same_Up | Transport reactions |
| MAR05087 | Same_Down | Same_Up | Same_Up | Transport reactions |
| MAR05088 | Same_Down | Same_Up | Same_Up | Transport reactions |
| MAR05089 | Same_Down | Same_Up | Same_Up | Transport reactions |
| MAR05305 | Same_Down | Same_Up | Same_Up | Transport reactions |
| MAR05308 | Same_Down | Same_Up | Same_Up | Transport reactions |
| MAR05320 | Same_Down | Same_Up | Same_Up | Transport reactions |
| MAR05552 | Same_Down | Same_Up | Same_Up | Transport reactions |
| MAR05578 | Same_Down | Same_Up | Same_Up | Transport reactions |
| MAR05583 | Same_Down | Same_Up | Same_Up | Transport reactions |
| MAR05586 | Same_Down | Same_Up | Same_Up | Transport reactions |
| MAR05587 | Same_Down | Same_Up | Same_Up | Transport reactions |
| MAR05588 | Same_Down | Same_Up | Same_Up | Transport reactions |
| MAR05590 | Same_Down | Same_Up | Same_Up | Transport reactions |
| MAR05593 | Same_Down | Same_Up | Same_Up | Transport reactions |
| MAR05594 | Same_Down | Same_Up | Same_Up | Transport reactions |
| MAR05595 | Same_Down | Same_Up | Same_Up | Transport reactions |
| MAR05596 | Same_Down | Same_Up | Same_Up | Transport reactions |
| MAR05597 | Same_Down | Same_Up | Same_Up | Transport reactions |

| | | | | |
|---|---|---|---|---|
| MAR05600 | Same_Down | Same_Up | Same_Up | Transport reactions |
| MAR05601 | Same_Down | Same_Up | Same_Up | Transport reactions |
| MAR05603 | Same_Down | Same_Up | Same_Up | Transport reactions |
| MAR05610 | Same_Down | Same_Up | Same_Up | Transport reactions |
| MAR05613 | Same_Down | Same_Up | Same_Up | Transport reactions |
| MAR05616 | Same_Down | Same_Up | Same_Up | Transport reactions |
| MAR05618 | Same_Down | Same_Up | Same_Up | Transport reactions |
| MAR05621 | Same_Down | Same_Up | Same_Up | Transport reactions |
| MAR05627 | Same_Down | Same_Up | Same_Up | Transport reactions |
| MAR05630 | Same_Down | Same_Up | Same_Up | Transport reactions |
| MAR05633 | Same_Down | Same_Up | Same_Up | Transport reactions |
| MAR05634 | Same_Down | Same_Up | Same_Up | Transport reactions |
| MAR05635 | Same_Down | Same_Up | Same_Up | Transport reactions |
| MAR05637 | Same_Down | Same_Up | Same_Up | Transport reactions |
| MAR05639 | Same_Down | Same_Up | Same_Up | Transport reactions |
| MAR05640 | Same_Down | Same_Up | Same_Up | Transport reactions |
| MAR05641 | Same_Down | Same_Up | Same_Up | Transport reactions |
| MAR05642 | Same_Down | Same_Up | Same_Up | Transport reactions |
| MAR05643 | Same_Down | Same_Up | Same_Up | Transport reactions |
| MAR05648 | Same_Down | Same_Up | Same_Up | Transport reactions |
| MAR05649 | Same_Down | Same_Up | Same_Up | Transport reactions |
| MAR05650 | Same_Down | Same_Up | Same_Up | Transport reactions |
| MAR05654 | Same_Down | Same_Up | Same_Up | Transport reactions |
| MAR05657 | Same_Down | Same_Up | Same_Up | Transport reactions |
| MAR05658 | Same_Down | Same_Up | Same_Up | Transport reactions |
| MAR05671 | Same_Down | Same_Up | Same_Up | Transport reactions |
| MAR05709 | Same_Down | Same_Up | Same_Up | Transport reactions |
| MAR05753 | Same_Down | Same_Up | Same_Up | Transport reactions |
| MAR05755 | Same_Down | Same_Up | Same_Up | Transport reactions |
| MAR05764 | Same_Down | Same_Up | Same_Up | Transport reactions |
| MAR05793 | Same_Down | Same_Up | Same_Up | Transport reactions |
| MAR06287 | Same_Down | Same_Up | Same_Up | Transport reactions |
| MAR06290 | Same_Down | Same_Up | Same_Up | Transport reactions |
| MAR06294 | Same_Down | Same_Up | Same_Up | Transport reactions |
| MAR06381 | Same_Down | Same_Up | Same_Up | Transport reactions |
| MAR06382 | Same_Down | Same_Up | Same_Up | Transport reactions |
| MAR07864 | Same_Down | Same_Up | Same_Up | Nucleotide metabolism |
| MAR07883 | Same_Down | Same_Up | Same_Up | Nucleotide metabolism |
| MAR07885 | Same_Down | Same_Up | Same_Up | Nucleotide metabolism |
| MAR08468 | Same_Down | Same_Up | Same_Up | Nucleotide metabolism |
| MAR08741 | Same_Down | Same_Up | Same_Up | Transport reactions |
| MAR20124 | Same_Down | Same_Up | Same_Up | Transport reactions |

Table S7: List of commonly perturbed reactions with same perturbation status in Bone and Brain (i.e., Down-regulated) but different in Liver (i.e., Up-regulated).

| **Reaction ID** | **Liver** | **Bone** | **Brain** | **Pathways** |
|---|---|---|---|---|
| MAR00155 | Same_Up | Same_Down | Same_Down | Transport reactions |
| MAR02303 | Same_Up | Same_Down | Same_Down | Transport reactions |
| MAR04171 | Same_Up | Same_Down | Same_Down | Purine metabolism |
| MAR05073 | Same_Up | Same_Down | Same_Down | Transport reactions |
| MAR05326 | Same_Up | Same_Down | Same_Down | Transport reactions |
| MAR05663 | Same_Up | Same_Down | Same_Down | Transport reactions |
| MAR05669 | Same_Up | Same_Down | Same_Down | Transport reactions |
| MAR05672 | Same_Up | Same_Down | Same_Down | Transport reactions |
| MAR06298 | Same_Up | Same_Down | Same_Down | Transport reactions |
| MAR07895 | Same_Up | Same_Down | Same_Down | Nucleotide metabolism |
| MAR08465 | Same_Up | Same_Down | Same_Down | Nucleotide metabolism |
| MAR08469 | Same_Up | Same_Down | Same_Down | Nucleotide metabolism |
| MAR08507 | Same_Up | Same_Down | Same_Down | Pyruvate metabolism |
| MAR08509 | Same_Up | Same_Down | Same_Down | Pyruvate metabolism |
| MAR20144 | Same_Up | Same_Down | Same_Down | Transport reactions |

Table S8: **Transport reactions with organ-specific regulation, associated metabolites, and genes:** This table summarizes transport reactions that show differential regulation (up- or down-regulation) across liver, bone, and brain tissues. For each reaction, the associated metabolites are listed, along with the genes encoding the corresponding transporters.

| **Reaction ID** | **Liver** | **Bone** | **Brain** | **Metabolites** | **Genes** |
|---|---|---|---|---|---|
| MAR05561 | Same_Up | Same_Up | Same_Down | MAM02136, MAM02360 | SLC7A5 |
| MAR05607 | | | | MAM01307, MAM02519, MAM02724 | SLC7A6, SLC7A5 |
| MAR05629 | | | | MAM01975, MAM02519, MAM02993 | SLC7A6, SLC1A5 |
| MAR05660 | | | | MAM02136, MAM02471, MAM02519 | SLC7A6, SLC7A5 |
| MAR05673 | | | | MAM02519, MAM03089, MAM03135 | SLC7A6, SLC7A5 |
| MAR05074 | Same_Down | Same_Down | Same_Up | MAM03101 | SLC7A2, SLC7A8, SLC7A5, SLC16A10, SLC36A1, SLC7A1, SLC43A1, SLC7A7, SLC7A3, SLC43A2, SLC3A2 |
| MAR05487 | | | | MAM01975, MAM02896 | SLC7A5, SLC7A10, SLC3A2 |
| MAR05659 | | | | MAM02471, MAM02519, MAM02993 | SLC7A6, SLC7A5 |
| MAR05674 | | | | MAM02519, MAM02993, MAM03089 | SLC7A6, SLC7A5 |
| MAR05580 | Same_Up | Same_Down | Same_Up | MAM02125, MAM02426 | SLC7A6, SLC7A5 |
| MAR05625 | | | | MAM01975, MAM02360, MAM02519 | SLC7A6, SLC7A5 |

| | | | | | |
|---|---|---|---|---|---|
| MAR05645 | Same_Down | Same_Up | Same_Down | MAM02136, MAM02519, MAM02896 | SLC7A6, SLC7A5 |
| MAR05664 | | | | MAM01975, MAM02519, MAM03089 | SLC7A6, SLC7A5 |
| MAR05873 | | | | MAM01365, MAM02360, MAM02519 | SLC7A6, SLC7A7, SLC3A2 |
| MAR05087 | Same_Down | Same_Up | Same_Up | MAM02770 | SLC7A2, SLC7A8, SLC7A5, SLC36A1, SLC7A1, SLC43A1, SLC7A7, SLC7A3, SLC43A2 |
| MAR05089 | | | | MAM03135 | SLC7A2, SLC7A8, SLC7A5, SLC36A1, SLC7A1, SLC43A1, SLC7A7, SLC7A3, SLC43A2 |
| MAR05583 | | | | MAM01365, MAM02125 | SLC7A6, SLC7A5 |
| MAR05586 | | | | MAM01588, MAM02125 | SLC7A6, SLC7A5 |
| MAR05587 | | | | MAM01307, MAM01986, MAM02519 | SLC7A6, SLC7A5 |
| MAR05588 | | | | MAM01975, MAM01986, MAM02519 | SLC7A6, SLC7A5 |
| MAR05590 | | | | MAM01986, MAM02471, MAM02519 | SLC7A6, SLC7A5 |
| MAR05593 | | | | MAM01986, MAM02519, MAM03101 | SLC7A6, SLC7A5 |

| | | | | | |
|---|---|---|---|---|---|
| MAR05594 | | | | MAM01628, MAM01986, MAM02519 | SLC7A6, SLC7A5 |
| MAR05595 | | | | MAM01986, MAM02360, MAM02519 | SLC7A6, SLC7A5 |
| MAR05596 | | | | MAM01986, MAM02519, MAM02770 | SLC7A6, SLC7A5 |
| MAR05597 | | | | MAM01369, MAM01986, MAM02519 | SLC7A6, SLC7A5 |
| MAR05600 | | | | MAM01986, MAM02136, MAM02519 | SLC7A6, SLC7A5 |
| MAR05601 | | | | MAM01986, MAM02184, MAM02519 | SLC7A6, SLC7A5 |
| MAR05603 | | | | MAM01307, MAM01975, MAM02519 | SLC7A6, SLC1A5 |
| MAR05610 | | | | MAM01307, MAM02360, MAM02519 | SLC7A6, SLC7A5 |
| MAR05613 | | | | MAM01307, MAM02519, MAM03135 | SLC7A6, SLC7A5 |
| MAR05616 | | | | MAM01307, MAM02184, MAM02519 | SLC7A6, SLC7A5 |
| MAR05618 | | | | MAM01307, MAM01975, MAM02519 | SLC7A6, SLC1A5 |
| MAR05621 | | | | MAM01975, MAM02519, MAM03089 | SLC7A6, SLC7A5 |
| MAR05627 | | | | MAM01369, MAM01975, MAM02519 | SLC7A6, SLC1A5 |
| MAR05630 | | | | MAM01975, MAM02136, MAM02519 | SLC7A6, SLC7A5 |
| MAR05633 | | | | MAM01307, MAM02519, MAM02896 | SLC7A6, SLC1A5, SLC1A4 |

| | | | | | |
|---|---|---|---|---|---|
| MAR05634 | | | | MAM01975, MAM02519, MAM02896 | SLC7A6, SLC1A5 |
| MAR05635 | | | | MAM02471, MAM02519, MAM02896 | SLC7A6, SLC7A5 |
| MAR05637 | | | | MAM02519, MAM02724, MAM02896 | SLC7A6, SLC7A5 |
| MAR05639 | | | | MAM01628, MAM02519, MAM02896 | SLC7A6, SLC1A5, SLC1A4 |
| MAR05640 | | | | MAM02360, MAM02519, MAM02896 | SLC7A6, SLC7A5 |
| MAR05641 | | | | MAM02519, MAM02770, MAM02896 | SLC7A6, SLC7A5 |
| MAR05642 | | | | MAM01369, MAM02519, MAM02896 | SLC7A6, SLC1A5 |
| MAR05643 | | | | MAM02519, MAM02896, MAM03135 | SLC7A6, SLC7A5 |
| MAR05648 | | | | MAM01307, MAM02471, MAM02519 | SLC7A6, SLC7A5 |
| MAR05649 | | | | MAM01975, MAM02471, MAM02519 | SLC7A6, SLC7A5 |
| MAR05650 | | | | MAM02471, MAM02519, MAM02896 | SLC7A6, SLC7A5 |
| MAR05654 | | | | MAM01628, MAM02471, MAM02519 | SLC7A6, SLC7A5 |
| MAR05657 | | | | MAM01369, MAM02471, MAM02519 | SLC7A6, SLC7A5 |
| MAR05658 | | | | MAM02471, MAM02519, MAM03135 | SLC7A6, SLC7A5 |
| MAR05671 | | | | MAM02519, MAM02770, MAM03089 | SLC7A6, SLC7A5 |

| | | | | | |
|---|---|---|---|---|---|
| MAR05709 | | | | MAM01628, MAM01975, MAM02519 | SLC7A6, SLC1A5, SLC1A4 |
| MAR05753 | | | | MAM01307, MAM01369, MAM02519 | SLC7A6, SLC1A5 |
| MAR05755 | | | | MAM01369, MAM02519, MAM02896 | SLC7A6, SLC1A5 |
| MAR05764 | | | | MAM01369, MAM02519, MAM02993 | SLC7A6, SLC1A5 |
| MAR05793 | | | | MAM01369, MAM02519, MAM02993 | SLC7A6, SLC1A5 |
| MAR06381 | | | | MAM02039, MAM02519, MAM02896 | SLC38A5, SLC38A1, SLC38A2, SLC38A4 |
| MAR06382 | | | | MAM01369, MAM02039, MAM02519 | SLC38A5, SLC38A1, SLC38A2, SLC38A4 |
| MAR05073 | Same_Up | Same_Down | Same_Down | MAM02471 | SLC7A2, SLC7A8, SLC7A5, SLC36A1, SLC7A1, SLC43A1, SLC7A7, SLC7A3, SLC43A2 |
| MAR05663 | | | | MAM01307, MAM02519, MAM03089 | SLC7A6, SLC7A5 |
| MAR05669 | | | | MAM01628, MAM02519, MAM03089 | SLC7A6, SLC7A5 |
| MAR05672 | | | | MAM01369, MAM02519, MAM03089 | SLC7A6, SLC7A5 |
